# The Emirati T2T-Level Pangenome: A Graph of 58 Complete Genomes

**DOI:** 10.1101/2024.12.16.628631

**Authors:** Michael Olbrich, Mira Mousa, Inken Wohlers, Amira Al Aamri, Halima Alnaqbi, Aisha Hanaya Alsuwaidi, Hima Vadakkeveettil Manoharan, Nour al-dain Marzouka, Sanjay Erathodi Ramachandran, Anju Annie Thomas, Mauricio Paton Gasso, Mubarak Almehairbi, Mohamed Alameri, Guan Tay, Rifat Hamoudi, Saleh Ibrahim, Noura Al Ghaithi, Habiba Alsafar

## Abstract

Reference data on genomic variation form the basis of genetic research. Limitations in identifying genetic variation from single reference sequences have been recently overcome, as improvements in sequencing technologies have allowed the generation of pangenomic references from multiple accurate, chromosome-level *de novo* assemblies. Here, we present a comprehensive Emirati telomere-to-telomere (T2T) pangenome generated from 58 individuals, comprising 28 trio-based and 30 single-sample assemblies. The resulting 116 haplotype-resolved assemblies demonstrate high contiguity, with a median continuity of 150 Mb and a median quality value (QV) of 59, achieving T2T-level scaffold status for 71.9% of chromosomes. These assemblies form the foundation of the Emirati T2T pangenome graph. The graph reveals levels of genomic diversity comparable to those reported by the Human Pangenome Reference Consortium, while also capturing regionally enriched and difficult-to-assemble variation, uniquely accessible through the Emirati T2T assemblies. This reference makes a valuable global contribution to human pangenomics and serves as a critical resource for advancing precision medicine in the United Arab Emirates.

## Introduction

The human reference genome serves as a foundational resource for genomics research and biomedical advancements, offering a common framework for interpreting human genetic variation. Landmark initiatives, such as the Human Pangenome Reference Consortium (HPRC)^1^ and the Telomere-to-Telomere (T2T) consortium^2^, have expanded this foundation. The HPRC introduced a draft pangenome comprising 47 phased, diploid assemblies from a cohort of genetically diverse individuals^1^. At the same time, the T2T consortium completed the first gapless, contiguous sequence of a haploid human genome, known as T2T-CHM13^2^. These milestones have provided open-access genomic resources critical for variant detection and interpretation, functional annotation, population genetics, and epigenetic analyses.

Despite these advances, current reference genomes capture only a narrow subset of global human diversity. The underrepresentation of many populations introduces reference bias, compromising the accuracy of variant calling and limiting insights into population-specific disease mechanisms. Efforts have increasingly turned toward the development of population-specific pangenomes that better reflect regional genetic landscapes. Projects such as the Chinese Pangenome^3^, the Arab Pangenome, and the African Pangenome have demonstrated the utility of these resources in enhancing variant discovery and informing population health. Despite these regional advancements, critical gaps remain in the representation of Middle Eastern populations, particularly those from the Arabian Peninsula. These populations possess distinct genetic features shaped by indigenous ancestry, high rates of consanguinity, and historical connectivity. Existing global and regional references have yet to fully capture this complexity, thereby limiting accurate variant interpretation and hindering the development of population-specific genomic tools.

The United Arab Emirates (UAE), through the Emirati Genome Program (EGP), is addressing this gap by undertaking one of the largest national genome initiatives, with over 700,000 samples collected to date from its citizen population of approximately one million. Here, we present the first telomere-to-telomere diploid Emirati pangenome, which incorporates a population-specific reference of exceptional completeness. This assembly resolves previously inaccessible genomic regions, providing a critical foundation for future studies. By capturing the unique genetic features of the Emirati population, this resource not only complements existing global pangenomes but also establishes a blueprint for embedding regional references into global genomic research frameworks.

## Results

### Cohort Selection for the Reference Assembly

The Emirati genome reference comprises 58 high-quality genome assemblies, including a female Emirati T2T assembly and 57 additional assemblies forming the Emirati pangenome dataset. These include 27 individuals from parent-offspring trios and 30 unrelated individuals. Individuals were deliberately chosen, building on the work of Mousa et al.^6^, to ensure representation of the UAE’s genetic diversity. The selection procedure adhered to the following criteria: (1) all individuals were selected from distinct population clusters stratified by the ancestral components of the Emirati population; the T2T reference sample was selected to represent the highest admixture proportion of the major ancestral component lineage; (2) all individuals were free of known congenital disorders, providing a healthy baseline for genomic analysis; and (3) trio families were restricted to individuals with unrelated parents, defined by a kinship coefficient no closer than the fourth degree, to ensure accurate genetic phasing. Fig. 1 illustrates the results of the population genetic admixture analysis and the sex distribution of the cohort, emphasizing the extensive genetic diversity captured within this dataset.

**Figure 1.**
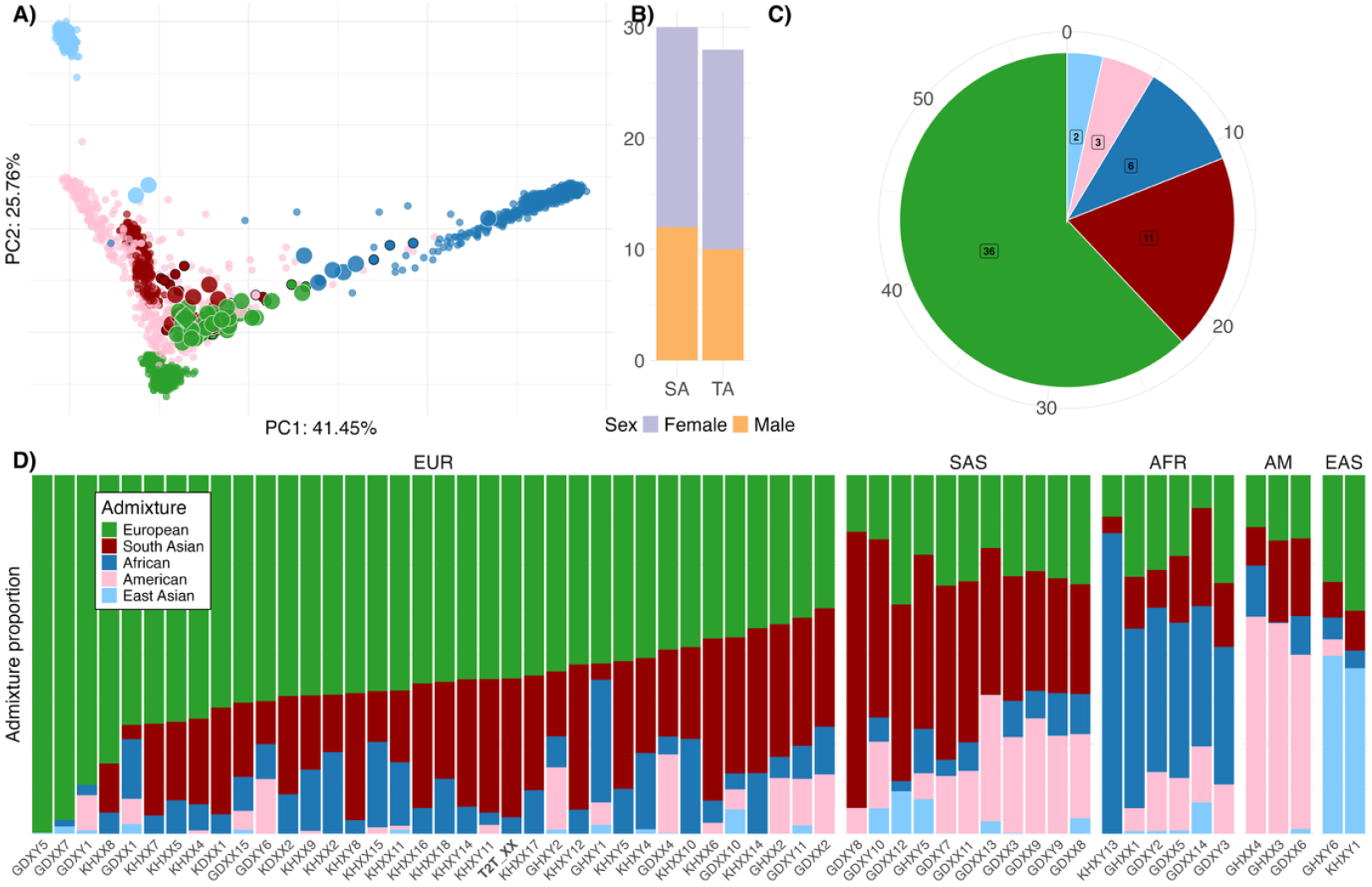
Basic and ancestry-based characteristics of the 58 genomes included in the Emirati genome reference. (a) The genomes are projected onto the genotype principal components computed by the 1000 Genomes phase 3 samples. Each Emirati genome is colored according to its 1000 Genomes-based global ancestry component with the maximum value. Larger dots with white borders highlight the assemblies, while in the case of trio assemblies, the parents are shown with smaller black-bordered dots. The T2T-ERG_XX sample is denoted by diamond-shaped symbols, with the T2T-ERG_XX genome located between the two parent genomes. (b) The number of pangenome single (SA) and trio (TA) assemblies stratified by sex, including the T2T assembly. (c) The distribution of the main ancestry components of the genomes across the cohort. (d) For each Emirati genome, the contributing ancestry components after supervised admixture analysis based on 1000 Genomes global populations, i.e., European, South Asian, African, American, and East Asian, c.f. Supplementary Table 17.

### Telomere-to-Telomere Assembly

A diploid, trio-based T2T assembly of a female Emirati individual using a combination of Illumina (Supplementary Table 1), PacBio HiFi (Supplementary Table 2), and ONT ultra-long sequencing data (Supplementary Table 2) was generated, referred to as T2T-ERG_XX in the following. The sequencing effort included 147.75X coverage of Illumina reads, 126.63X coverage from PacBio HiFi for the child (Supplementary Table 2), and 64.05X and 63.01X coverage of PacBio HiFi reads for the mother and father, respectively (Supplementary Table 3). The data were complemented by 111.8X coverage from ONT ultra-long reads, generated from 19 ONT PromethION sequencing runs, with a median N50 of 67.8 kb (Supplementary Table 2). PacBio HiFi data were derived from four sequencing runs, producing a median N50 of 17.9 kb and a coverage of 126.6X.

The assembly approach, as outlined in Supplementary Fig. 1, involved multiple stages of read trimming, polishing, scaffolding, and assembly refinement using a range of tools. The quality of the assemblies was evaluated at various stages using a balance of contiguity (number of contigs and auN/N50 Values), base accuracy (QV score), and completeness (percentage of N values) (Supplementary Table 4). The final assembly was selected based on its optimal trade-off between these parameters and is derived from an initial assembly with Verkko^7^, scaffolding with ntLink^8^, followed by CHM13 reference-guided scaffolding using RagTag^9^, and gap-filling via two alternative hifiasm^10^ assemblies using quarTeT^11^ (Supplementary Fig. 1).

The final assembly comprises chromosome-level haplotypes with an overall length of 3.03 Gb. The maternal haplotype has an adjusted QV score of 60.8, and the paternal haplotype has an adjusted QV score of 59.8, with switch errors of 2% and 3.2%, respectively (Supplementary Table 5). The assembly contains 101 gaps, of which 37 are in the maternal haplotype and 64 in the paternal haplotype (Supplementary Table 6). The corresponding N-bases per 100 kb are 2.21 for the maternal and 7.10 for the paternal haplotype (Supplementary Table 5). Notably, chromosomes 4, 6, 12, 14, 15, 18, 21, and 22 in the maternal haplotype and chromosomes 6 and 8 in the paternal haplotype are gap-free (Supplementary Table 6), indicating that these chromosomes were fully assembled from telomere to telomere.

The mitochondrial genome was not incorporated during assembly but was reconstructed using Illumina short-read-based mitochondrial variant calls and aligned to the rCRS. Our assembly covered 94.4% of the CHM13 sequence, with 32.5 and 29.2 Mb of the maternal and paternal haplotypes, respectively, not aligning to CHM13, corresponding to approximately 1% of the genome. The total CHM13 aligned length was 2.99 Gb.

In terms of gene coverage, our assembly included 96.6% of the genes reported in CHM13 (62,060 of 64,000 genes), with 258 genes detected only partially. Of the 1,550 multi-copy genes in CHM13, 1,219 were also present with multiple copies in our assembly (78.6%), amounting to 4,746 genes (Supplementary Table 5). Interestingly, these numbers were identical for both the maternal and paternal haplotypes. After manually fixing a misjoin between chromosomes 14 and 15 (also reported and corrected for CHM13), five and four interchromosomal misjoins in the maternal and paternal haplotypes, respectively, were identified when compared to CHM13.

Alignment of our assembly with CHM13 showed that all chromosomes are assembled accurately (Fig. 2a maternal and Supplementary Fig. 2a paternal haplotype) but revealed that centromeric regions were partially unaligned. A few larger centromeric regions in chr13, chr14, chr15, chr18, chr21, and chr22 aligned between non-homologous chromosomes, highlighting differences that relate to assembly challenges that were reported for CHM13 previously (Fig. 2b-c and Supplementary Fig. 2b-c). Reads from chromosome X were also aligned to the pseudoautosomal regions of the Y chromosome in CHM13, despite our assembly not containing a Y chromosome.

**Figure 2.**
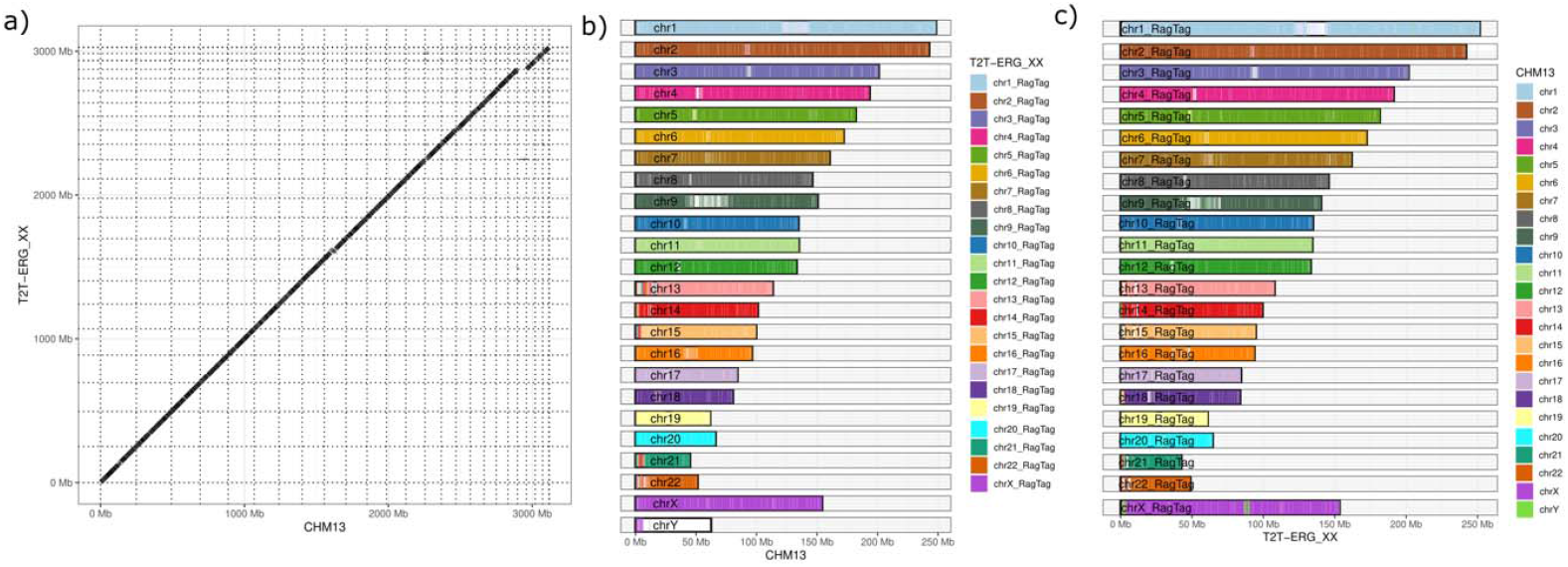
Alignment of T2T-ERG_XX maternal haplotype with CHM13v2.0. All primary alignments are shown. (a) Dotplot. (b) The coverage of T2T-ERG_XX chromosomes with CHM13 chromosomes. (c) The coverage of CHM13 chromosomes with T2T-ERG_XX chromosomes. In (b) and (c), all assembly sequences with a length of more than 5 Mb are shown.

### Fifty-Seven High-Quality Diploid Assemblies

The T2T assembly of the genome of an Emirati individual was complemented with the assemblies of 57 representative genomes. The underlying data was generated by one PacBio Revio Run per sample, with ten samples receiving an additional sequencing top-up to enhance coverage. The median coverage per sample was 31.8X, with a range of 25.5X to 52.4X, and a median read N50 of 16.8 kb (Supplementary Table 7 and Supplementary Fig. 3). Adapter trimming was performed using HiFi data, but no adapters were detected, and contamination screening was conducted with Kraken2 ^12^. Assembly was performed using hifiasm ^10^, considering parental HiFi data for the 27 trio assemblies to generate fully phased diploid haplotypes. Scaffolding was then performed using ntLink ^8^, followed by CHM13 reference-guided scaffolding using RagTag ^9^, and subsequent error correction and polishing with Inspector ^13^ (Supplementary Fig. 4 summarizes the pangenome processing strategy).

The assembly quality and characteristics were evaluated at the draft hifiasm haplotypes stage (Supplementary Table 8) and the final haplotype stage (Supplementary Table 9 and Supplementary Fig. 5), with quality improving through post-processing. For the final assemblies’ haplotypes, the median number of contigs was 193, with a range of 100 to 335 (see Fig. 3a). The average N50 was 150.32 Mb, with a range of 140.92 MB to 154.94 Mb (see Fig. 3b) Our assemblies showed cumulative contig length curves that aligned closely with CHM13, reflecting similar assembly continuity and indicating high assembly quality (see Fig. 3c). The median overall length across all haplotypes was 3.02 Gb. For the 70 haplotypes from female genomes (which contain no Y chromosome and diploid X chromosomes), the median length was 3.02 Gb. The paternal haplotypes of ten male individuals from trio assemblies had a median length of 2.93 Gb (see Fig. 3d), confirming accurate phasing of diploid genomes without an X chromosome.

**Figure 3.**
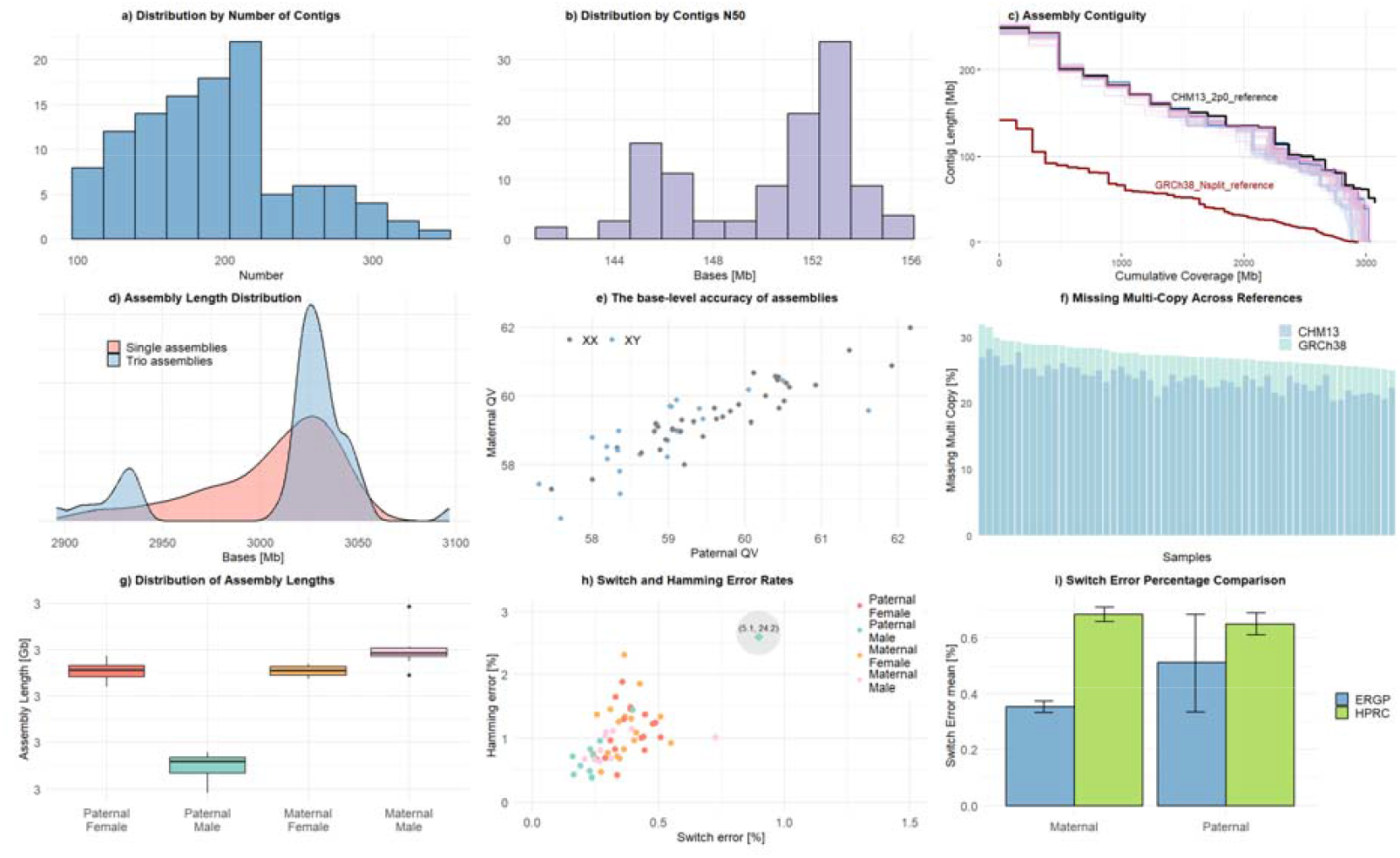
Assembly characteristics of 57 post-processed HiFi assemblies. Panel (a) presents the distribution by the number of contigs for all assemblies. Panel (b) shows the N50 length across assemblies. Panel (c) shows the contig length vs. cumulative assembly coverage, compared with CHM13 and GRCh38 references. Panel (d) displays the overall sequence length for 114 haplotypes, stratified by single-sample and trio-assembly approaches. Panel (e) compares the quality values (QV) between paternal and maternal haplotypes. Panel (f) illustrates the percentage of missing multi-copy genes. Panel (g) summarizes total contig lengths stratified by sex and haplotype for trio-assembly samples. Panel (h) plots Hamming error versus switch error (%) for the 27 trio assemblies. Note: the highlighted outlier corresponds to a male paternal haplotype with elevated Hamming error; see the main text for details. Panel (i) compares the mean switch error of all haplotypes from the 27 trio assemblies with the mean switch error reported by the Human Pangenome Reference Consortium (HPRC).

The adjusted QV score of maternal and paternal haplotypes was highly correlated, with values ranging from 56.44 to 62.15, and a median of 59.31 (Fig. 3e). Analysis of the interchromosomal misjoins showed a median of 3 misjoins in the range of 0 to 7 misjoins for all haplotypes. When accounting for the complexity of mapping centromeric regions, no interchromosomal misjoins were reported. A median of 61,541 of the 64,214 genes reported in CHM13 were detected across the haplotypes, representing 95.83% of the total. A median of 360 genes was detected only partially (Supplementary Table 9). Of the 1,550 multi-copy genes reported in CHM13, a median of 1,181 (76%) were detected in our assembly.

When comparing our assemblies to the GRCh38 reference, the percentage of missing multi-copy genes was higher and less variable across haplotypes compared to the CHM13 reference, as shown in Fig. 3f. We also observed differences in haplotype lengths for the male genome, with the paternal haplotype being shorter due to the presence of the Y chromosome, instead of an X chromosome. Male maternal haplotypes tended to be longer than female maternal haplotypes, although both covered the same chromosomes (Fig. 3g). The length of the manually curated MT genomes ranges from 16,566 to 16,574 bases (median 16,569 bases), matching the GRCh38 reference length of 16,569 bases.

Switch and Hamming errors were assessed across the 54 haplotypes from trio assemblies and were affected by sex chromosomes. Switch errors across all haplotypes were low, with a median of 0.33% (average: 0.43%, [0.16, 5.05]), with a single haplotype showing a switch error rate of 5.05% (Fig. 3h), closely outperforming the switch error rate of the HPRC pangenome assemblies (Fig. 3i). The Hamming error showed a median of 0.9% (avg: 1.4%, [0.3, 24.1]) over all assemblies. The male assemblies showed the lowest switch error (median 0.2%) and Hamming error (median 0.7%) rates (Fig. 3h), with the notable exception of one male paternal assembly, which showed a Hamming error of 24.14% and a switch error of 5.05%.

### Emirati Pangenomic Variation

The Minigraph-Cactus^14^ pipeline was employed to generate two distinct pangenome graphs. For each graph, either CHM13v2 or GRCh38 served as the primary backbone, complemented by the alternative reference genome included as an unclipped additional sequence, followed by the inclusion of the two T2T-ERG_XX haplotypes and the 114 haplotypes from the 57 HiFi-based assemblies (Supplementary Fig. 6). The number of nodes, edges, and corresponding sequence lengths for each graph configuration is summarized in Supplementary Table 10.

The CHM13v2-backboned graph consistently displayed higher values across all major metrics. It contained 121.3 million nodes, 88.4 million nodes, and spanned 3.35 Gb, compared to 111.3 million edges, 79.4 million nodes, and 3.23 Gb in the GRCh38-backboned graph. These differences were especially pronounced on larger chromosomes. For example, chromosome 1 in the CHM13v2-backboned graph reached around 9.3 million edges, 6.76 million nodes, and 267.4 Mb, while the GRCh38-based counterpart had around 8.1 million edges, 5.86 million nodes, and 263.9 Mb.

Across nearly all autosomes, CHM13v2-based graphs exhibited longer graph sequence lengths and higher node counts. This trend was especially marked on chromosomes 13 through 22, which contain complex and repetitive regions better resolved in CHM13. For instance, chromosome 13 in the CHM13 graph included around 3.03 million nodes and spanned 121.2 Mb, while in the GRCh38 graph, it was reduced to around 2.37 million nodes and 117.2 Mb.

Sex chromosomes also showed large differences, especially chrY, which exhibited 1.9-fold more edges in the CHM13-based graph (4.4 million vs. 2.3 million) and almost double the node count (3.29 million vs. 1.74 million), demonstrating the more complete representation of this chromosome in the CHM13-based assembly. Conversely, chrX showed only minor differences in graph length and topology between the two backbones.

The total added sequence from the 58 Emirati individuals that CHM13 and GRCh38 do not capture was 222.69 Mb (Supplementary Table 11), with the per-genome contribution detailed in Fig. 4a. The inclusion of T2T-ERG_XX haplotypes contributed 1.46 Mb of novel sequence over the CHM13v3 and GRCh38 reference backbone. Furthermore, 566,159 novel bases, shared by at least 55 Emirati individuals, were identified as the core graph. When adding genomes to the pangenome, the novel sequence added per genome constitutes increasingly only sequence private to the respective genome (Supplementary Fig. 7).

**Figure 4.**
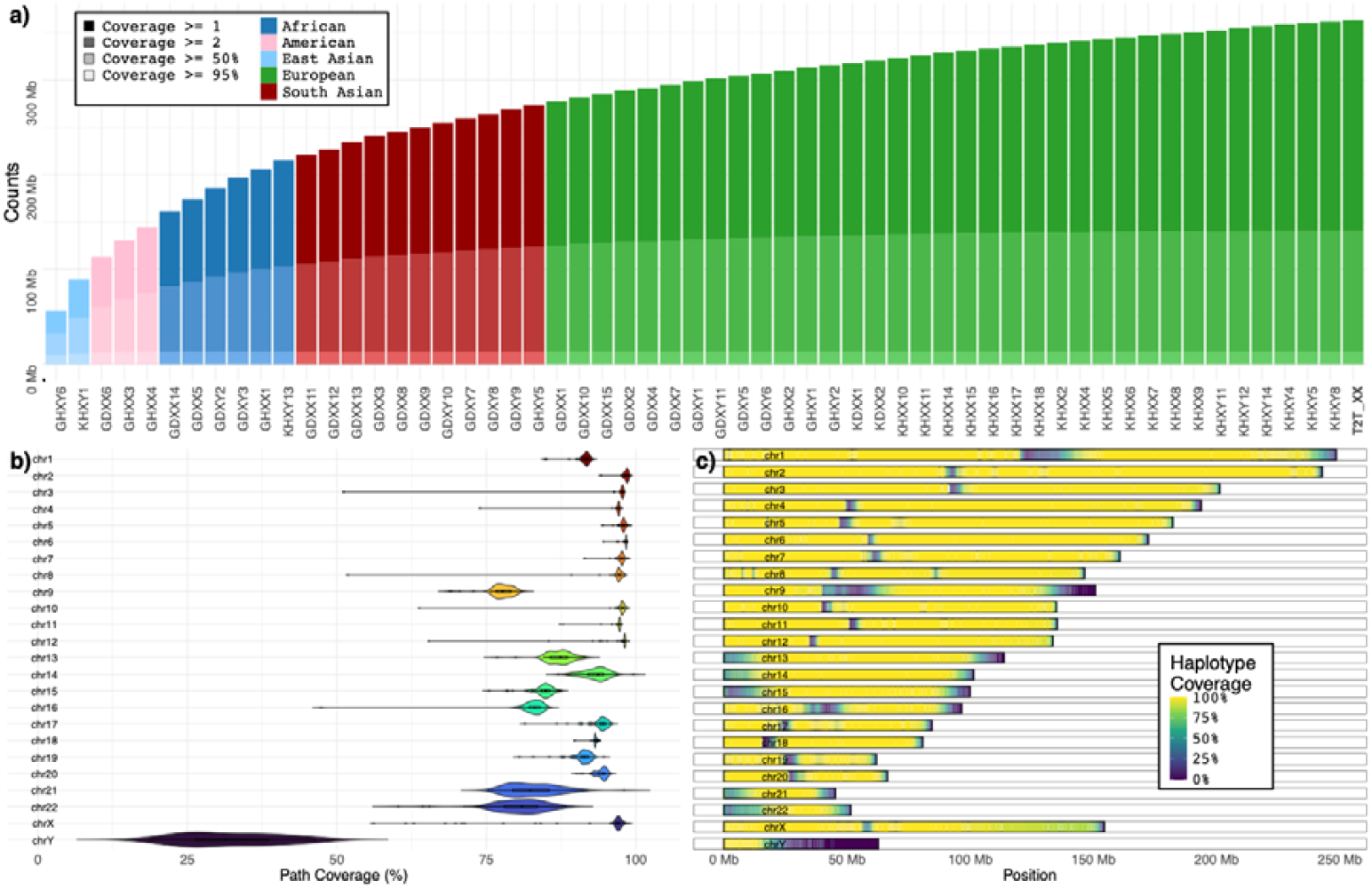
Characteristics of the pangenome graph (a) Bar plots depict the ordered growth of the pangenome graph in base pairs across successive sample additions. Values are reported for individual sample contributions as well as cumulative growth when samples are added in Panacus order, from 1 to 58. The growth curve demonstrates a steep initial increase in graph size with the first samples, followed by a gradual plateau, indicating diminishing contribution to the total graph as additional genomes are incorporated. (b) Violin plot of graph segment coverage across chromosomes in 116 scaffold assemblies mapped to the CHM13 reference. The median segment coverage for all autosomes ranges from 77.7% (chromosome 9) to 98.5% (chromosome 2). Coverage exceeded 97% for most large autosomes (chromosomes 2–8, 10–12), while lower median values were observed for chromosomes 9 (77.7%), 22 (80.9%), 16 (83.1%), 21 (82.3%), and 15 (84.9%). Median path coverage across sex chromosomes. The median coverage was 97.0% for chromosome X and 31.4% for chromosome Y. (c) A heatmap of path coverage across the backbone of CHM13 chromosomes, where each position is colored by the fraction of assembled haplotype contigs that traverse the graph at that locus. The pattern recapitulates the distribution summarized in panel (b), with high coverage along most autosomes and chromosome X, and pronounced depletion in repetitive regions. Coverage is lowest on the chromosome Y, consistent with its median of 31.4%, and remains high on the chromosome X, with a median of 97.0%. Reduced signal is concentrated in centromeric and subtelomeric intervals and on the short arms of the acrocentric chromosomes 13, 14, 15, 21, and 22, reflecting expected challenges in assembling highly repetitive sequences.

Variant counts were calculated compared to GRCh38 and CHM13 in the two graph configurations, respectively. Variant assessment across these graphs revealed substantial differences in variant representation depending on the selected backbone (Supplementary Table 12; Supplementary Fig. 8). The CHM13v2-backboned graph consistently captured the highest number of total variants, with 22,995,968 variant records when using CHM13 as the reference and 20,720,149 records when using GRCh38. In contrast, the GRCh38-backboned graph had 21,287,028 records when integrating CHM13 and 21,584,845 records with GRCh38 alone. The CHM13-backboned graph using CHM13 also showed the most comprehensive capture of single-nucleotide variants (SNVs), with 19,232,359 SNVs, compared to 17,893,818–18,019,495 SNPs in the GRCh38-backboned graph. This trend was consistent across other variant types. CHM13-based graphs had more multiallelic sites (up to 1,601,705) and multiallelic SNV sites (277,978) than the GRCh38-backboned counterparts. Notably, the number of multinucleated polymorphisms (MNPs) was markedly higher in the CHM13v2-backboned graph with CHM13 as reference (1,016,706 MNPs) compared to GRCh38 (462,591) or the GRCh38-backboned graphs (ranging from 744,065 to 923,243). Indels followed a similar trend, with the CHM13-based graph detecting 3,138,525 indels, the highest across all conditions. Between 98.6 and 99.8% of variant records are SNVs or insertions and deletions (summarized as indels), with their overall number in relation to the three references in the Emirati T2T pangenome graph shown in Supplementary Figure 8.

The pangenome graph paths relating to the CHM13 chromosomes (Supplementary Table 13), i.e., the proportion of the CHM13 reference chromosome path covered by the longest graph traversal per haplotype, for each of the 57 reference samples and the independent sample T2T-ERG_XX were scrutinized (Supplementary Table 14, Supplementary Table 15). The coverage across autosomes, sex chromosomes, and the mitochondrial chromosome (chrM), stratified by haplotype, was assessed. Across the 57-sample pangenome graph, autosomal CHM13 path coverage was consistently high (Figure 4b), with a median of 94.53% of CHM13 chromosome covered. Notably, chromosome 21 exhibited the most compact range of segment coverage (median 99.4%), while chromosomes 13 and 15 had slightly lower coverage distributions, driven by variability in contig continuity and alignment gaps.

In contrast, coverage across sex chromosomes showed greater variability. For chrX, coverage was generally high in XX haplotypes (median 98.4%), but lower in XY individuals, where chrX coverage was limited to the non-pseudoautosomal regions (median 86.2%). ChrY coverage was substantially lower and more heterogeneous, with many samples missing coverage altogether or displaying highly fragmented shared graph sequence (median 65.7% among those with nonzero coverage). These differences reflect both biological variation and technical challenges in representing repetitive Y-chromosome sequences within a graph framework. ChrM (mitochondrial DNA) exhibited near-complete coverage in most haplotypes, with a median mitochondrial path coverage exceeding 95%.

Primary contigs that contributed meaningfully to chromosome-level coverage were lifted over to CHM13v2 and evaluated for both coverage and gap burden. In aggregate, these contigs mapped 3.37 × 10^11 base pairs to the reference. Reference coverage was high, with a mean of 91.74% and a median of 94.89%. Of the 2,679 contigs, 2,584 contained at least one N-defined scaffold gap (Supplementary Table 16), and 95 were gap-free, including 57 manually curated mitochondrial contigs. The combined gap content across all assemblies and all contigs was 0.001152108%. Overall, there were only three autosomal CHM13 regions that were not covered by any of the Emirati assembly graph paths, a 5.6 kb centromeric region on chromosome 1, a 130 kb centromeric satellite region on chromosome 11 and a 1.25 Mb region on chromosome 13 (Supplementary Table 17).

N-region sizes of unknown bases show a clear mode at 100 base pairs, consistent with the scaffolding stitching procedure, and a long tail of larger events revealed by the axis break (Supplementary Figure 9a). A higher total N content per haplotype chromosome was associated with a lower percent coverage of the CHM13 pangenome graph path (Supplementary Figure 9b), and this inverse trend was evident after excluding sex chromosomes. Together, these findings indicate high overall coverage of the reference by assembled contigs, widespread but low-level scaffold gaps that cluster near centromeres and telomeres, and a measurable impact of aggregate gap burden on effective CHM13 coverage.

Focusing on the T2T-ERG_XX sample, we observed uniformly high segment coverage across both haplotypes for all autosomes, with values ranging from 97.7% to 99.9%. ChrX coverage was similarly robust in both haplotypes (99.3% and 99.7%), consistent with an XX karyotype. ChrY segments were absent, and chrM coverage was complete in both haplotypes (100%), suggesting comprehensive mitochondrial representation.

The number of SNVs per genome identified through the pangenome assembly graph (Supplementary Figure 8) was compared to those obtained from traditional read mapping-based variant calling using Sentieon, based on the same HiFi data that was used for assembly. Per genome SNV counts from mapping-based calls ranged from 4.08 million to 5.68 million, with a median of 4.59 million (Supplementary Table 18). Genomes with predominant African ancestry (Supplementary Table 19) consistently showed the highest SNV counts, exceeding 5 million in all five African individuals, and reaching a maximum of 5.68 million SNVs in a single genome (KHXY13). One European main component genome also approached this range, with 5.05 million SNVs.

In contrast, SNV counts derived from the pangenome graph were uniformly lower. When aligned to a CHM13 backbone, the median SNV count across the 57 samples was 3.66 million. Using GRCh38 as a backbone yielded slightly higher SNV counts, with a median of 3.70 million, representing a 1.09% increase compared to CHM13. Individual differences in SNV counts between the two backbones reached up to 2.19%, corresponding to a maximum difference of 87,842 SNVs. The reference genome T2T-ERG_XX yielded 3.75 million SNVs using a GRCh38 backbone and 3.66 million using CHM13, which closely aligns with the corresponding population medians (Supplementary Figure 10). In general, African main component genomes retained the highest counts across all pangenome graph configurations. For CHM13 backbone-based graphs, African main component genomes exhibited between 4.07 and 4.38 million SNVs, compared to 3.38 to 3.79 million for the majority of European main component genomes. East Asian and South Asian main component genomes showed intermediate values. These results underscore that ancestry influences both the total number of SNVs and the degree to which they are captured by graph-based versus linear reference-based approaches.

### Locus-Specific Graph-Based Analysis of Complex HLA Structural Variants

To contextualize our findings, we performed a locus-specific analysis of clinically relevant complex SV loci within the HLA in the Emirati pangenome using an approach comparable to that described in the HPRC publication. Like the HPRC, we used graph-based assemblies to resolve large multi-allelic structural variants and to assess their representation relative to linear references (GRCh38, CHM13).

Fig. 5a shows the graph representation of clinically relevant complex SV loci within the HLA region constructed from the current Emirati T2T pangenome. We identified five looped structures (L1–L5) corresponding to large-scale insertion–deletion polymorphisms (≥50 bp to 44 kb in range). These alternative paths reflect distinct structural haplotypes spanning key loci of 10 genes (HCP5B, HLA-U, HLA-K, HLA-A, HLA-H, HLA-Y, HLA-W, HLA-J, HCG4B, and HCG9).

**Figure 5:**
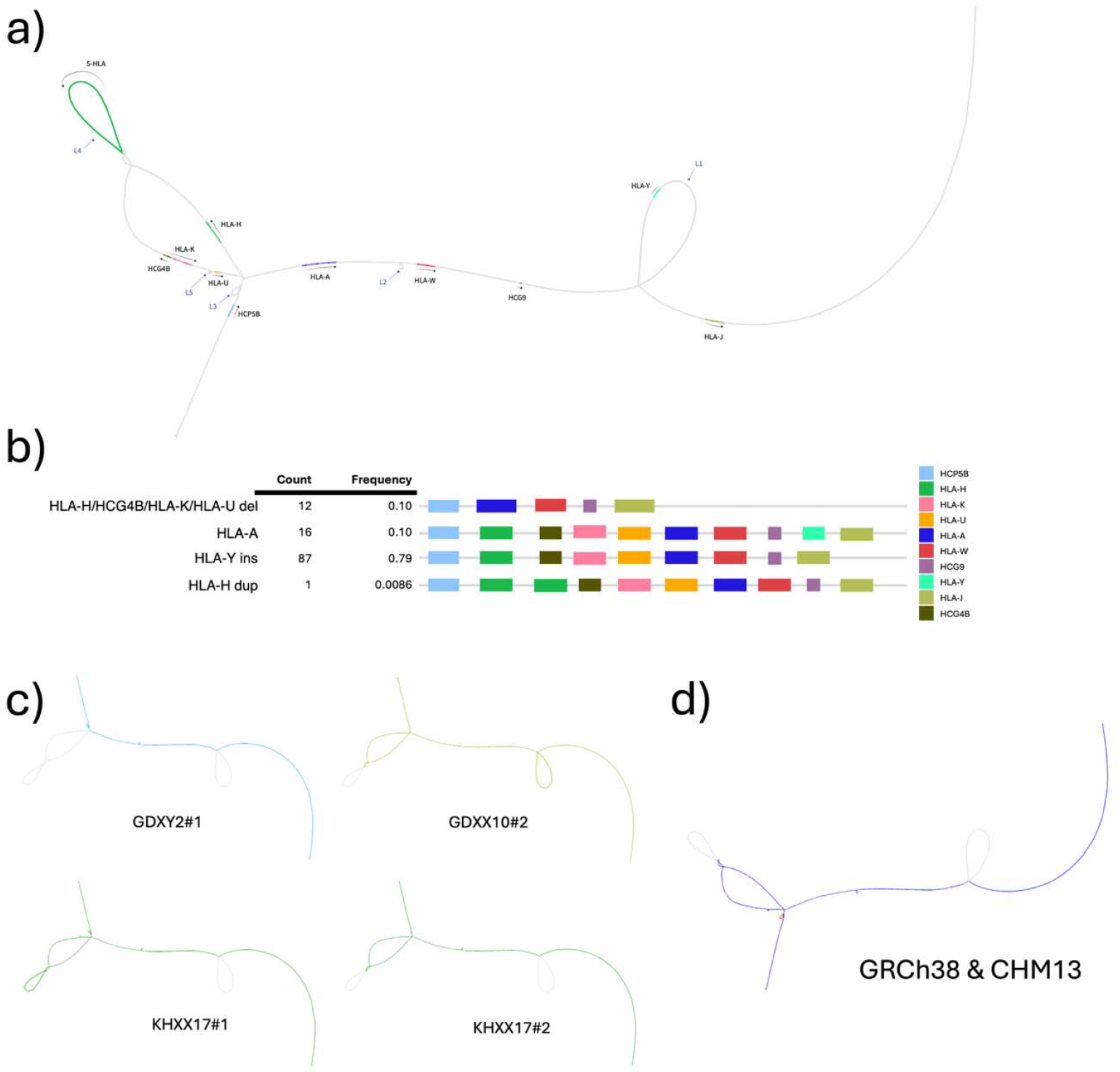
Visualizing complex pangenome loci. (a) Structural variation of HLA-A from the MC graph. b) Haplotype frequencies across the analyzed assemblies c-d) Representative haplotypes illustrate distinct structural configurations relative to GRCh38 & CHM13v2, including HLA-A, HLA-Y insertion, HLA-H/HCG4B/HLA-K/HLA-U deletion, and HLA-H duplication.

Notably, L1 represents a 65.8 kb insertion carrying the HLA-Y pseudogene, while L4 corresponds to a 44.0 kb alternative path containing HLA-H. The presence of these large loops indicates reference-absent haplotypes not represented in GRCh38 or CHM13v2. Additional loops (L2, L3, L5) encompass smaller structural variants that overlap with canonical HLA and neighbouring genes. The HLA-Y insertion (L1) is supported by 16 haplotypes (13.8%), with one individual (GDXX10) carrying the insertion on both chromosomes. The multi-gene deletion spanning HLA-H, HCG4B, HLA-K, and HLA-U occurs on 12 haplotypes (10.3%).

In contrast, the L4 haplotype (HLA-H insertion) in Fig. 5c is observed only once (KHXX17) (0.86%). We also detected a multi-gene deletion spanning HLA-H, HCG4B, HLA-K, and HLA-U in 12 haplotypes (∼10.3%), including two individuals (KDXX1, KHXY1) who were homozygous for the deletion. Stratification by parental haplotype revealed modest asymmetry, with maternal haplotypes contributing more frequently to both the HLA-Y insertion (11 maternal vs. 5 paternal) and the multi-gene deletion (8 maternal vs. 4 paternal) (Fig. 5; Supplementary Table 1).

Our findings mirror and extend those of the HPRC. Consistently, we observe the approximately 65 kb HLA-Y insertion, although at a lower frequency (∼14% in Emiratis vs. ∼28% in HPRC, globally). Importantly, our analysis reveals additional structural complexity not captured in HPRC datasets, including a rare homozygous HLA-H haplotype in (L4).

### Emirati Reference-Based Variant Detection

An independent cohort of 119 Emirati genomes was used for graph evaluation. It was initially characterized with respect to basic demographic and ancestry features (Supplementary Fig. 11). Short⍰read alignment and variant detection across these genomes (median depth 42X, range 31–77X) were assessed against five reference configurations: the linear GRCh38 and T2T⍰CHM13 assemblies, the HPRC pangenome graph constructed on an hg38 backbone, and two Emirati pangenome graph configurations described earlier, one rooted in hg38 and the other in CHM13 (Supplementary Fig. 12), and the intersection of variant calls investigated (Supplementary Fig. 13).

GRCh38 remained the most permissive target for BWA⍰MEM, with a median of 93.6% of reads and 94.7% of bases mapped (Supplementary Table 21). CHM13 matched GRCh38 within 0.1%, while the three graph-based references (HPRC and the two graphs generated as part of this work) mapped approximately one percentage point fewer reads. Despite this small decrease in raw mappability, graph alignment, or simply replacing the hg38 backbone with CHM13, reduced duplicate alignments by a median of 1.5million reads per sample (approximately 2%), so the unique depth remained well above 30X in every genome.

Relative to linear GRCh38, graph alignment yielded a median increase of 1.9% in single⍰nucleotide variants (≈80,000 additional SNVs) and 1.8% in small insertions or deletions (≈18,000 additional indels) per genome. The uplift persisted outside RepeatMasker regions (≈1% more variants outside of repeat regions), indicating genuine sensitivity rather than repeat⍰driven inflation. CHM13 alone recovered approximately half of the graph advantage. In contrast, tailoring the graph to the Emirati cohort (Emirati T2T pangenome versus the global HPRC graph) altered total SNV and indel counts by less than 0.2%. The two graphs alternated in their per-sample ranking, suggesting that a high-quality generic graph already captures most short-read diversity in this population.

A different trend was observed for insertions and deletions larger than 50 bp outside annotated repeats. Alignment to linear GRCh38 followed by structural variant calling produced a median of 165,000 insertions or deletions longer than 50 bases outside annotated repeats, whereas both Emirati T2T pangenome graphs, the HPRC graph, and linear CHM13 suppressed these events by ∼96% (to fewer than 10,000 per sample). This reduction indicates that most insertion and deletion structural variations reported against GRCh38 are alignment artefacts that disappear when reads are mapped to a graph or the more complete CHM13 assembly.

In summary, replacing linear GRCh38 with a modern backbone, either graph⍰based or CHM13, delivers approximately 2% higher small⍰variant sensitivity, a 2% reduction in duplicates, and near⍰elimination of spurious insertion or deletion structural variants, while incurring only a one⍰percentage⍰point loss in raw mappability.

#### Benchmarking Analysis of HG001-HG007 Samples

The seven Genome-in-a-Bottle (GIAB) genomes (HG001 to HG007), with both short and long reads, were analyzed by aligning each dataset to three references: the linear GRCh38 assembly and both the HPRC and Emirati T2T pangenome graphs with a GRCh38 backbone (Supplementary Fig. 14; Supplementary Table 22). Small variant calling was performed and benchmarked against the GIAB truth sets for single-nucleotide polymorphisms (SNPs) (Supplementary Figs. 15, 16, 17) and small insertions or deletions (indels) (Supplementary Figs. 18, 19, 20).

SNV detection was effectively saturated across all references and technologies. With long reads, median F1 scores were 0.9993 on GRCh38, 0.9983 on the HPRC graph, and 0.9979 on the Emirati T2T pangenome graph; short reads yielded 0.9956, 0.9950, and 0.9948, respectively. The largest absolute gap between any graph and the linear assembly was 0.00014. GRCh38 also produced the fewest false positives, for example, 7,115 versus 10,403 on the HPRC graph and 12,483 on the Emirati T2T pangenome graph in HG001 short reads. These results indicate that transitioning from a linear to a graph-based reference does not materially impact SNV calling accuracy.

Reference choice influenced indel performance more strongly and interacted with read length. Long reads on GRCh38 achieved a median F1 score of 0.9937, compared with 0.9803 on the HPRC graph and 0.9770 on the Emirati T2T pangenome graph, reflecting roughly 1,500 additional true positives and up to 2,000 fewer false positives per genome. In HG005, long⍰read indel F1 score was 0.9857 on GRCh38, 0.9722 on the HPRC graph, and 0.9667 on the Emirati graph. With short reads, all three references converged: median F1 score clustered between 0.9928 and 0.9931, with graphs sometimes recovering a few hundred extra variants at a modest cost in precision.

Read length exerted the greatest overall influence. Moving from short to long reads improved the single⍰nucleotide polymorphism F1 score by approximately 0.003 on the linear reference and 0.002 on each graph. For indels, the gain approached two percentage points and reduced false negatives by an order of magnitude, as in HG003, where the count fell from 3,764 to 256 on GRCh38.

Error profiles differed by variant class. Graph-based alignment introduced 1.5-3 times more SNV false positives than linear assembly. It led to slightly lower transition-to-transversion in long-read samples, with 1.9987 for linear GRCh38 and 1.84 for both graph references. The heterozygous-to-homozygous ratios for both long and short read samples, with 1.5633 for linear short read, 1.49 for graph-based short reads, and 1.68 for linear long reads and 1.58 for graph-based long reads. For indels, long reads on GRCh38 reduced both false negatives and false positives, whereas short reads on graphs traded a slight rise in precision for a similar rise in recall.

Overall, GRCh38 paired with long reads delivered the cleanest call sets, but graph references closed much of the gap for indels with short reads and never lagged by more than 0.003 F1 score in SNV detection. The Emirati T2T pangenome graph performed within 0.3% of the HPRC graph across all metrics, suggesting a limited additional benefit from population-specific tailoring for well-characterized genomes.

## Discussion

This study presents the first complete high-quality, telomere-to-telomere diploid pangenome from the Emirati population, addressing the historical underrepresentation of the Middle Eastern population in global genomics initiatives. Our cohort was systematically selected based on admixture profiles, ensuring comprehensive representation, and trio-based assemblies with unrelated parents were included to support reliable haplotype phasing.

The diploid, trio-based T2T assembly of a female Emirati individual’s genome achieves 94.4% coverage of the CHM13 sequence, with 96.6% of the genes represented, including multi-copy genes, and gap-free maternal haplotypes for chromosomes 4, 6, 12, 14, 15, 18, 21, and 22. The maternal and paternal haplotypes were well-phased, with minor discrepancies in centromeric regions and interchromosomal misjoins affecting previously reported regions, which are common challenges in genome assembly due to their repetitive nature. Notably, our assembly also revealed structural variations, particularly in the centromeric regions, highlighting the genomic diversity within the Emirati population.

Extending to 57 additional haplotype-resolved assemblies, we observed median contig N50 of 150.32 Mb and a median QV of 59, with 71.9% of chromosomes achieving T2T completeness. Inclusion of all 58 genomes in the pangenome graph added approximately 222.69 Mb of novel sequence not represented in CHM13 or GRCh38, with a core of 566 kb shared by at least 55 individuals. This highlights population-specific features that are absent from existing global references, including multiple structural variants and novel sequences, potentially relevant for gene regulation or disease susceptibility.

Compared with HPRC, the Chinese Pangenome consortium (CPC), and Arab reference efforts, our assemblies demonstrate contiguity, base-calling accuracy (QV 59.31) and phasing fidelity (switch error rate 0.47%). In terms of assembly contiguity, our average N50 of 150.32 Mb surpasses those of the HPRC (40 Mb), CPC (>35.63 MB), and Arab (124.25 Mb) references, highlighting the strength of our assembly pipeline. Notably, our N50 was achieved with an average sequencing coverage (30.9X HiFi). Similarly, our QV score of 59.31 exceeds those of the HPRC (53.57), CPC (48–60), and Arab reference (57.53), indicating a high degree of base-calling accuracy and assembly fidelity. Furthermore, our switch error rate of 0.47% is the lowest among the pangenomes evaluated, outperforming the HPRC (0.67%), CPC, and Arab reference (0.64%), and demonstrating superior phasing and haplotype accuracy.

The graph’s variant record count in relation to the maternal T2T-EGR_XX haplotype in our Emirati pangenome was 20.84 million, falling between the variant counts observed for GRCh38 and CHM13, the previous human genome references. Individual genome’s SNV counts in the pangenome graph ranged from 4.08 million to 5.68 million, with a median of 4.59 million, showing significant diversity, particularly in genomes with tAfrican main global ancestry components. In contrast, the pangenome graph itself showed a reduction in SNV counts, with a median of 3.66 million SNVs when aligned to the CHM13 backbone, representing a 3.89% reduction compared to the CHM13 reference. These results align closely with previous findings in pangenome studies, where the genomic variation of different populations has been mapped to reference genomes with varying degrees of resolution.

These results emphasize the value of regional pangenomes in capturing rare, ancestry-specific alleles that can be underrepresented or absent in global reference datasets. Within the HLA region, such alleles have profound implications because of their direct relevance to clinical genetics and population health. By resolving structural haplotypes that are invisible in GRCh38 and CHM13, the Emirati T2T pangenome reveals variants that are important for refining our understanding of immune-related disease susceptibility and for improving the reliability of pharmacogenomic risk prediction. Importantly, the discovery of population-specific haplotypes (such as the ∼44 kb HLA-H allele) and the recurrent multi-gene deletion demonstrates that ancestry-tailored genomic resources are indispensable for ensuring equity in precision medicine.

We acknowledge important limitations. Structural variants represented by single or very few haplotypes (such as the ∼44 kb HLA-H path L4) may reflect true but ultra-rare alleles, yet they could also represent assembly artifacts, phasing errors, or graph misalignments. Similarly, the detection of large multi-gene deletions must be interpreted cautiously, as highly repetitive sequence content and paralogous gene families in the HLA region increase the risk of spurious structural reconstructions.

These uncertainties underscore the importance of orthogonal validation, including long-read resequencing and targeted PCR, before attributing functional or clinical significance to novel alleles.

Moving forward, several steps are critical: (i) expanding sample size and ancestry representation will help distinguish true population-specific variants from assembly noise. (ii) Integration of functional genomic assays can determine whether novel haplotypes exert any biological effects. (iii) developing robust benchmarking frameworks for complex loci to ensure reproducibility across graph-construction pipelines. Ultimately, the integration of regional and global pangenome efforts will be crucial to strike a balance between population-specific resolution and global comparability.

Our evaluation of both global and Emirati pangenome graphs using Genome in a Bottle (GIAB) benchmarks and an independent cohort of 119 Emirati genomes underscores the growing utility of graph-based and comprehensive references in short-read variant discovery. While SNV detection is now effectively saturated across technologies and references, indel calling remains more sensitive to both the underlying reference and the sequencing platform. As expected, long-read data aligned to GRCh38 yielded the most precise indel callsets, reflecting extensive optimization against benchmark datasets. However, for short-read data, both the HPRC and Emirati graphs enhanced the recall of small indels and effectively suppressed spurious large insertions and deletions, with only modest increases in false positives and minimal reductions in mappability. The linear CHM13 reference achieved comparable improvements, indicating that reference completeness alone can substantially improve alignment and variant detection.

The incorporation of Emirati haplotypes into a population-specific graph yielded minimal additional benefit when benchmarking against GIAB samples, likely due to the absence of population-matched variation. However, it lays essential groundwork for improved sensitivity in underrepresented populations. In the Emirati cohort, where no validated truth set exists, graph-based methods consistently recovered a subset of variants missed by linear alignment, even across samples with varying sequence quality. The high concordance across references suggests robust variant detection overall, yet the distinct variants uniquely captured by graph approaches highlight the importance of continued methodological refinement. Collectively, these results support a hybrid strategy: graph-based alignment for comprehensive discovery, and GRCh38 with long reads for high-confidence validation. The benchmarking framework employed here, including both global references and a newly constructed population-specific graph, provides a rigorous assessment of real-world performance. As graph mapping and variant calling methods continue to mature, particularly for long reads, their integration will help close existing gaps and support more accurate, equitable genomic analysis across diverse populations.

Our study advances equitable, population-specific genomics by providing the first telomere-to-telomere diploid pangenome for the UAE. By capturing the unique genetic architecture of the Emirati population, our pangenome graph not only closes a significant gap in the worldwide representation of human diversity but also enables more accurate and inclusive genomic analyses. This resource supports precision medicine in the UAE and broader Middle Eastern populations, where high rates of consanguinity and unique population structure make tailored references critical for rare diseases, carrier screening, and pharmacogenomic stratification. Its incorporation into national genomic initiatives, such as the Emirati Genome Program, promises to reduce diagnostic delays and support the integration of genomic data into clinical decision-making. Beyond its regional significance, our approach, including high-coverage long reads, trio-based phasing, T2T assembly, and pangenome graph construction, offers a scalable framework for other populations, while benchmarking against GIAB and Emirati samples informs accurate variant calling, particularly for indels and complex regions.

The Emirati T2T pangenome presents a foundational resource for clinical genomics, facilitating more accurate interpretation of functional variants and supporting the development of diagnostic and therapeutic strategies tailored to the population. Beyond its national relevance, it offers a methodological framework for integrating regional genomic diversity into global reference efforts, advancing the goal of inclusive and equitable precision medicine.

## Methods

### Sample Selection

Parent-child trios and single samples were identified from the Emirati Genome Program (EGP) and local recruitment sites to capture the genetic diversity of the UAE population^6^. The EGP dataset was initially screened for congenital disorders to exclude affected individuals, thereby ensuring that the cohort consisted of genetically unaffected individuals. Familial trios were then identified, with a focus on selecting unrelated parents, as this distance in kinship (up to the fourth degree) represents the algorithmic cutoff for reliable genomic analysis.

To assess familial relationships, KING^15^ software was used to infer relatedness up to the fourth degree, employing algorithms for kinship estimation and identity-by-descent (IBD) segment analysis. Following this, a graph-based approach in R was applied to visualize and further refine familial relationships. The analysis began with the construction of a subgraph focusing on parent-offspring relationships, enabling the identification of familial clusters. Clusters containing fewer than three individuals were excluded, and adjustments were made to account for varying cluster sizes. Additionally, these clusters were examined for consanguinity, with parent relationships inferred up to the third degree. Sex assignments, derived from DRAGEN^16^ ploidy information, were used to verify the correct parent-child assignments.

Due to the complexity of classifying familial relationships, certain groups were excluded in specific cases, including: (1) parents with first- or second-degree relationships; (2) full-sibling relationships, particularly when a trio involved two brothers and the offspring of one brother; and (3) cases involving full- or half-sibling relationships, where trios resembled consanguineous relationships, such as a child born between two half-siblings. Additionally, multigenerational configurations, such as a trio consisting of a grandfather, father, and grandson, were excluded due to ambiguities in kinship, such as a “half-sibling edge” between the grandfather and grandson, which complicated the accurate classification of parent-offspring connections.

### Admixture Analysis

Given the genetic diversity of the UAE population, a population genetic admixture analysis using Admixture^17^ was performed to stratify the population into distinct genetic subgroups. This stratification formed the basis for selecting familial trios that accurately represented the population’s genetic diversity. To further enhance the representation, a complementary sampling approach was employed to include single-sample assemblies alongside the familial trios. These single samples underwent the same admixture stratification to ensure their genetic background was consistent with the broader population structure.

A set of global population-informative variants from 12 studies, curated by Wohlers *et al*.^18^, was utilized. This dataset encompassed 127,261 variants observed in 5,429 individuals across 144 diverse populations selected for admixture analyses. This cohort was stratified according to continental origins based on five global populations of 3,502 individuals as part of the 1000 Genomes Project^19^. Each of the five components can be considered to have distinct global genetic ancestry, namely, European, South Asian, African, South American, and East Asian ancestry. Employing a supervised mode of admixture, the proportions of genetic ancestry contributed by these five distinct admixture components to the genomes of the Emirati population were quantified. This analysis allowed us to determine ancestral ratios, revealing the underlying global genetic substructure of the Emirati population. Additionally, principal component analysis (PCA) was performed using Plink^20^ on population-specific variants from the reference samples. The PCA enabled the projection of the Emirati population within the context of these principal components, facilitating ordination within the broader genetic framework of the global reference populations.

### Sequencing

#### Illumina Sequencing

The genomic DNA was extracted from 200 µL of EDTA-whole blood using QIAamp DNA Blood Mini Kit (Qiagen, Germany) according to the manufacturer’s instructions. The quality of the extracted genomic DNA (gDNA) was examined using a Nanodrop 1000 spectrophotometer (Thermo Fisher Scientific, USA) to determine A260/230 and A260/280 ratios. The integrity of the gDNA was evaluated through gel electrophoresis methods, while samples were quantified using the Qubit dsDNA BR assay kit on Qubit 4.0 Fluorometer (Thermo Fisher Scientific, USA).

Subsequently, the gDNA samples were normalized to 200 ng in 30 µL and proceeded with library preparation using Illumina DNA PCR-Free Library Prep kit and IDT® for Illumina® DNA/RNA UD Indexes Set A for library barcoding. Libraries were quantified using KAPA library quantification assay (KAPA Biosystems, USA) on ViiA 7 qPCR System (Applied Biosystems, USA), were normalized to 1.1 nM, and then pooled at equal volumes, followed by denaturation and further dilution before machine loading and sequencing. The S4 flowcells generated 150 bp paired end reads in a standard workflow. Short-read sequencing of the final libraries was carried out using S4 300 cycles Reagent Kit v1.5 on a NovaSeq 6000 Sequencer (Illumina, USA).

A 30X read coverage was estimated for the average human genome size (∼3.2 Gb) to perform sequencing on the NovaSeq 6000 (Illumina, USA) platform. All the samples were sequenced to achieve 96 Gb of raw data (FASTQ), corresponding to 30X coverage of the whole genome. Some tolerance was allowed, and WGS with a yield of >87 Gb was accepted. Samples falling short of the 96 Gb, and 30X autosomal sequencing coverage threshold underwent an additional step of top-up that included repeating the library preparation and barcoding. The number of samples selected for pooling in the top-up process was determined by the additional data required and the throughput capacity of the NovaSeq 6000 S4 Reagent Kit v1.5 (300 cycles). Following sequencing, all sample BCL files were generated and demultiplexed utilizing bcl2fastq on the DRAGEN platform (Illumina).

#### High-Fidelity (HiFi) Sequencing

The PacBio Nanobind CBB kit (Pacific Biosciences, 102-301-900) was used to extract high-molecular-weight (HMW) DNA from 200 µL peripheral whole blood samples. This extraction yields DNA fragments ranging from 50 to 500+ Kb, making it ideal for HiFi long-read WGS sequencing on the PacBio system. The concentration of the extracted gDNA samples was examined using the Qubit dsDNA BR kit fluorometric assay on Qubit 4.0 Fluorometer (Thermo Fisher Scientific, USA), and the fragment size was assessed using the gDNA 165 Kb kit on a Femto Pulse System (Agilent Technologies, USA). All DNA samples achieved the recommended Genome Quality Number (GQN) of 9.0 or above at 10 kb and 5.0 or above at 30 kb. The samples were then processed with a PacBio Short Read Eliminator (SRE) kit, following the manufacturer’s guidelines, to remove DNA fragments under 10 kb in length selectively. The concentration and GQN of the gDNA were determined using the Qubit 1x dsDNA HS kit fluorometric assay and the Femto Pulse - gDNA 165Kb method, respectively. Subsequently, sample shearing was executed using the Megaruptor 3 system (Diagenode s.a, Belgium) at a speed of 29-31 to achieve target insert lengths of 15-18 Kb, followed by sample cleanup using 1.0 X SMRTbell cleanup beads. The concentration and size distribution of the fragmented DNA were assessed using the Qubit 1x dsDNA HS kit fluorometric method and the Femto Pulse - gDNA 165Kb assay, respectively. Libraries of 12-20 kb size were then constructed using the PacBio SMRTbell prep kit 3.0 protocol (Pacific Biosciences, USA). Library preparation was performed using the manufacturer’s protocol, including DNA repair, A-tailing, adaptor ligation, and cleanup, followed by nuclease treatment to eliminate unligated DNA fragments and residual SMRTbell adapters. AMPure PB beads with a 3.1X ratio of beads (35%) were used to remove the shorter fragments (<5 kb) and concentrate the DNA sample. The concentration of the SMRTbell libraries was assessed using the Qubit 1x dsDNA HS fluorometric assay, followed by the Femto Pulse - gDNA 165Kb method to determine the fragment size distribution.

SMRTLink (v12.0) Sample Setup software was used to obtain instructions for WGS HiFi Reads Sequencing preparation (ABC), including primer annealing and polymerase binding. The final libraries were sequenced on a single 8 M SMRT Cell on the PacBio Revio platform.

#### Oxford Nanopore (ONT) Ultra-long Sequencing

To extract ultra-high molecular weight (UHMW) DNA for Oxford nanopore ultra-long sequencing, the buffy coat from a fresh 5-ml blood sample tube was aliquoted for peripheral blood mononuclear cells (PBMCs) isolation using RBC Lysis Solution (QIAGEN, 158904). The purified PBMCs were counted, and approximately 60 million cells per sample were carried forward for UHMW DNA extraction using the NEB Monarch Cells & Blood DNA extraction kit (New England Biolabs, T3050), with modifications as recommended by the manufacturer for ONT Ultra long sequencing application. The gDNA was eluted in the ONT Extraction EB (EEB) buffer.

The extracted gDNA was quantified using a method developed by Paul A ‘Giron’ Koetsier & Eric J Cantor, 2021. Subsequently, 30 to 40 μg of DNA in 750 μl of EEB buffer was used in the library preparation step using the ONT Ultra-Long DNA Sequencing Kit V14 (SQK-ULK114) in accordance with the supplied protocol. The library preparation began with an initial tagmentation step at room temperature for 10 minutes, followed by 10 minutes at 75°C, and then kept on ice for an additional 10 minutes. The rapid adapters were added and incubated for 30 minutes at room temperature; then, the adapter-ligated DNA was cleaned using a precipitation method. The final libraries were quantified, and 90 μL of the library was mixed with the sequencing buffer and the loading solution and kept at RT for 30 min. To proceed with the loading, the flow cells were primed with the priming mix as per protocol. After adding the libraries, they were incubated in the flow cell for 10 minutes. The sequencing setup was done using the MinKNOW software. The pores were scanned every 2 hours for optimal loading, and live base calling was conducted using the high-accuracy (HAC) base-calling model with base detection for 5mC on the MinKNOW software. The flow cells were reloaded two more times following the flow cell wash cycle as described in the protocol.

### Pangenome Graph Construction

The construction of the pangenome graph complemented with T2T-ERG_XX assembly followed a systematic, multi-step approach to ensure high data quality and accuracy. Supplementary Fig. 6 depicts this process, which involved assembling 30 single samples using the standard hifiasm^10^ mode, the phased assembly of 27 family trios in ‘trio-dual’ mode, and the generation of a primary T2T assembly, T2T-ERG_XX using a dedicated strategy. Details of the assembly strategies are provided in the respective sections. The graph was constructed using the current version of the Minigraph-Cactus^14^ pipeline. Two distinct pangenome graphs were generated to evaluate the Emirati pangenome. These graphs incorporated CHM13 as the primary backbone and GRCh38 as an additional unclipped reference and vice versa, with additionally 116 haplotypes from the 58 Emirati genomes, including the T2T-ERG_XX assembly. This graph was subsequently filtered to keep only nodes covered by more than 12 haplotypes (>20%) to obtain the Emirati T2T pangenome reference.

### Telomere-to-Telomere Assembly

The T2T-ERG_XX assembly strategy is outlined in Supplementary Fig. 1. Briefly, the process began with the removal of adapters from the HiFi data using HiFiAdapterFilt^21^, with no adapters detected, and from ONT data using Porechop_ABI^22^. ONT reads longer than 20 kb were filtered and then combined with PacBio HiFi data to construct a diploid, trio-based draft assembly using Verkko^7^. Parental k-mers, generated from the HiFi data with Meryl^23^, were employed in this process. The initial draft assembly was then scaffolded using ntLink^8^, followed by reference-guided scaffolding using RagTag^9^ with the CHM13 version 2 reference genome. To address gaps in the assembly, gap-filling was performed using two alternative hifiasm-based assemblies. These assemblies were generated from yak^24^-preprocessed parental HiFi data. The first hifiasm assembly was constructed using the same ONT and HiFi data as the Verkko assembly. For the second assembly, the ONT reads were polished with Ratatosk^25^, using Illumina data that had been downsampled to 50% with BBMap^26^, before performing the assembly. Finally, contigs from both hifiasm draft assemblies were utilized to polish the scaffolded Verkko assembly using QuarTeT^11^, resulting in the final assembly. Key assembly quality measures for selecting the optimal assembly strategy at different stages, as well as for the selected alternative assembly strategies, are provided in Supplementary Table 4. Further assembly characteristics of the final assembly can be found in Supplementary Table 5.

Alignment of T2T-EGR_XX with CHM13 was performed using minimap2^27^ alignment followed by filtering to keep primary alignments and visualization using the R package pafR. Dot plots and coverage plots were generated for all primary alignments. Coverage plots display only contigs with length more than 5 Mb.

### Pangenome Assembly

The core pangenome assemblies constructed using PacBio HiFi sequencing data comprise 28 phased trio assemblies (TAs) and 30 single assemblies (SAs). Single-sample assemblies were generated using hifiasm^10^ in standard mode, while trio-based assemblies were phased using hifiasm’s trio-dual mode. Sequencing data were preprocessed to remove adapters using HiFiAdapterFilt^21^, which confirmed the absence of adapter sequences across all datasets. Parental k-mer databases were created using yak^24^ with recommended settings for trio assemblies to ensure precise phasing and haplotype assignment.

Initial quality control assessments of the generated assemblies were conducted using Quast^28^ and yak^24^. Post-assembly scaffolding was performed with ntLink^8^ to optimize assembly continuity. Kraken2^12^ screening was carried out against human and non-human reference databases to identify and remove potential contamination. Chromosomal misjoins were detected using ‘paftools misjoin’, leveraging the CHM13v2 reference genome. This process identified three interchromosomal misjoins in the maternal haplotypes of samples GDXX5, GHXX3, and GDXX9, specifically: a misjoin between chromosomes 1 and 3 in GDXX5 (ntLink_4), chromosomes 10 and 12 in GHXX3 (ntLink_15), and chromosomes 1 and 13 in GDXX9 (ntLink_10). Each misjoin was resolved by splitting the affected contigs at the midpoint (with padding of 2 to 1,500 bp) and annotating the resulting fragments with suffixes’_part1’ and ‘_part2’ to maintain traceability.

Gene duplication analysis was conducted using ‘paftools asmgene’ against GRCh38 and CHM13v2 references, revealing no significant outliers. Further error correction at the base pair and structural variant levels was performed using Inspector, utilizing the original HiFi sequencing data as input.

A systematic renaming scheme was introduced to maintain consistency with pangenome standards. Each sample was encoded with identifiers reflecting its source (Genome Project or extended sampling), assembly type (haploid or diploid), inferred sex (XX or XY), and a sequential numerical ID. Contig names were updated to include the new sample ID and haplotype information, enabling seamless integration into the pangenome graph.

Manual curation of contigs was performed with reference to the mitochondrial genome by excluding contigs that match mitochondrial sequence but do not have the expected length, followed by linearizing the MT contigs to reflect GRCh38 reference positions. Therefore, we obtain a bona fide *de novo* assembled mitochondrial genome sequence for all assemblies.

Finally, secondary quality control checks using Quast^28^ and yak^24^ confirmed the accuracy of the polished assemblies.

### Pangenome Graph Assessment

The structural properties and genomic content of the pangenome graphs were evaluated using Panacus^29^, a tool specifically designed to assess pangenome graph structure. Key graph metrics were computed for each configuration, including the number of nodes, edges, and sequence lengths. To further characterize the graphs, the contribution of novel sequences from the T2T-ERG_XX assembly and the 57 Emirati genomes was quantified. Additionally, the overlap of graph paths with CHM13v2.0 and GRCh38 chromosomes was measured, allowing for an evaluation of coverage and single nucleotide and structural variation representation.

Variant calling was conducted using both linear reference-based and graph-based approaches. For linear references, Sentieon^30^’s variant-calling pipeline processed short-read Illumina sequencing data aligned to CHM13 and GRCh38, generating detailed variant statistics, including counts of single-nucleotide variants (SNVs) and small insertions and deletions (indels). This workflow ensured a consistent framework for comparing variants detected through graph-based mapping.

The VG Giraffe^31^ aligner maps short-read sequencing data to the constructed pangenome graphs. This alignment enabled Sentieon to call variants directly from the pangenome, producing variant records relative to the CHM13 backbone. Additionally, the variant call outputs from the Minigraph-Cactus^14^ pipeline were integrated into the analysis to evaluate graph-derived variants comprehensively. Finally, the sequence coverage of CHM13 chromosome paths by graph paths was measured to assess the contiguity of scaffolds within the pangenome.

For annotation of the HLA regions, the reference sequences were obtained from the IPD-IMGT/HLA database and Ensembl release v114. These regions were incorporated into the analysis by alignment of the selected sequences using GraphAligner to the Minigraph-cactus graph. The resulting GFA graph file was chunked to isolate the HLA region with a 20,000 base pair threshold, and the final graph was visualized using Bandage.

### Read-mapping based Evaluations

To evaluate the performance of graph-based and linear reference genomes for variant calling, 119 whole-genome sequencing (WGS) short-read samples previously published by Daw Elbait et al. (2021)^32^ were analyzed. Our comparison included two local graph references (Emirati T2T pangenome graphs: *ku_hg38_d12* and *ku_chm13_d12*), two public linear references (hg38 and CHM13), and a public graph reference (*HPRC_hg38_d9*). FASTQ files underwent quality control using FastQC, and alignment was performed using either linear or graph-based strategies. Reads were aligned to the GRCh38 or CHM13 reference genomes using the Sentieon DNAseq pipeline^30^. Alignment to the graphs was performed using the Variation Graph (VG) toolkit^31^ and variant calling was subsequently performed with Sentieon^30^ to ensure consistency across both approaches. Post-alignment processing and variant filtration were performed using SAMtools^33^ for BAM and CRAM files and BCFtools for VCF files containing single-nucleotide variants (SNVs), small insertions and deletions (indels), and structural variants (SVs). Several quality metrics across references were computed and compared, including the total number of SNPs, the number of homozygous and heterozygous SNPs, the number of indels, and the number of mapped bases. Results were visualized using custom plotting scripts. Additionally, the QC assessments were repeated after masking problematic repeat-rich genomic regions to evaluate the robustness of each reference system in complex regions.

To further benchmark variant calling accuracy, the performance was evaluated using Genome in a Bottle (GIAB)^34^ truth sets for seven widely used reference samples (HG001 to HG007). These samples comprise individuals of European, Ashkenazi Jewish, and Han Chinese ancestry, with established high-confidence variant calls and corresponding BED files that define regions suitable for benchmarking.

Publicly available Illumina short-read and PacBio HiFi long-read sequencing data were analyzed. Alignment to the linear GRCh38 reference was conducted using the Sentieon^30^ pipeline, and graph-based alignment was performed using the VG toolkit, with variant calling again executed through Sentieon^30^. To assess the accuracy of the called variants, hap.py (v0.3.14, Illumina)^35^ with default parameters and the supplied high-confidence BED file were employed.

## Supporting information

Main Figures

Supplementary Figures and Descriptions

## Data Availability

This study used the publicly available Genome in a Bottle (GIAB) benchmark dataset. Short-read sequencing data for GIAB samples HG001–HG007 were obtained from the GIAB NCBI FTP repository (short reads: https://ftp-trace.ncbi.nlm.nih.gov/ReferenceSamples/giab/data/). Long-read sequencing data for HG001–HG007 were obtained from the Human Pangenome/NHGRI–UCSC panel S3 bucket (long reads: https://s3-us-west-2.amazonaws.com/human-pangenomics/index.html?prefix=NHGRI_UCSC_panel/). Corresponding GIAB truth sets were downloaded from the GIAB release repository (https://ftp-trace.ncbi.nlm.nih.gov/ReferenceSamples/giab/release/).

Reference resources included CHM13 (from MARBL; https://github.com/marbl/CHM13), GRCh38 (Ensembl release 113; https://ftp.ensembl.org/pub/release-113/fasta/homo_sapiens/dna), and Human Pangenome graph resources (https://github.com/human-pangenomics/hpp_pangenome_resources). New data generated here, Emirati T2T assemblies, trio assemblies, single-individual assemblies, and the Emirati pangenome graph have been submitted to the European Genome-phenome Archive (EGA). They will be available under controlled access upon publication under EGA accessions: Study ID [EGAS50000001232 - EGAS50000001235]; Dataset ID(s) [EGAD50000001753 - EGAD50000001756]; Sample/Analysis ID(s) [EGAZ50000006419 - EGAZ50000006422].

## Code Availability

The list of all tools, including versions and/or code commits, used for this study are available in Supplementary Table 23.

## Acknowledgements

We would like to extend our sincere gratitude to all the individuals who generously donated their samples to the EGP and to the local recruitment sites. Their willingness to participate in this study is invaluable and forms the foundation of this research, contributing to our understanding of the genetic landscape of the UAE population. Special thanks to Salsabeel A.S. Juneidi for recruitment and DNA extraction, Sarah Chehadeh for implementation of the ONT-UL protocol, and Khacey Juma for recruitment. Their expertise and dedication have been instrumental in ensuring the project’s success.

## Author Contributions

HS, SI, MM, and NA designed the study. MO, IW, and HVM developed the assembly strategy, with MO and HVM carrying out T2T assembly construction. MO, AA, and AHA constructed the pangenome assemblies, while AA led the post-processing and QC of the assemblies. MO, IW, AA, and HVM examined the characteristics of the assemblies. MO and AHA were responsible for creating the pangenome graph, while AHA led the post-processing and visualization of the pangenome, with IW and AHA analyzing its characteristics. MO, MM, HAN, and AA handled data coordination and management. NaM, MA, and MPG conducted the graph variant calling evaluation. The original draft was written by MO and MM, with IW contributing, while MA, GT, RH, SA, and NaG undertook the review and editing of the manuscript. Admixture analysis was conducted by MO and NaM. Sample selection was carried out by MO and MM. HAN, SAAS, AAT, and SER performed sequencing. HS served as the principal investigator and oversaw the laboratory organization that HAN managed. The overall conceptualization, funding acquisition, and supervision of the project were led by HS. All authors reviewed and approved the final draft of the manuscript.

## Ethics Declarations

The study procedures and protocols adhered to the ethical guidelines and regulations established by the Research Ethics Scientific Committee in the UAE (DOH: DOH/CVDC/2022/1701) and Khalifa University (KU: H21-045), as well as the principles outlined in the Declaration of Helsinki.

