## Supplementary material for "The Emirati T2T-Level Pangenome: A Graph of 58 Complete Genomes": Main Figures

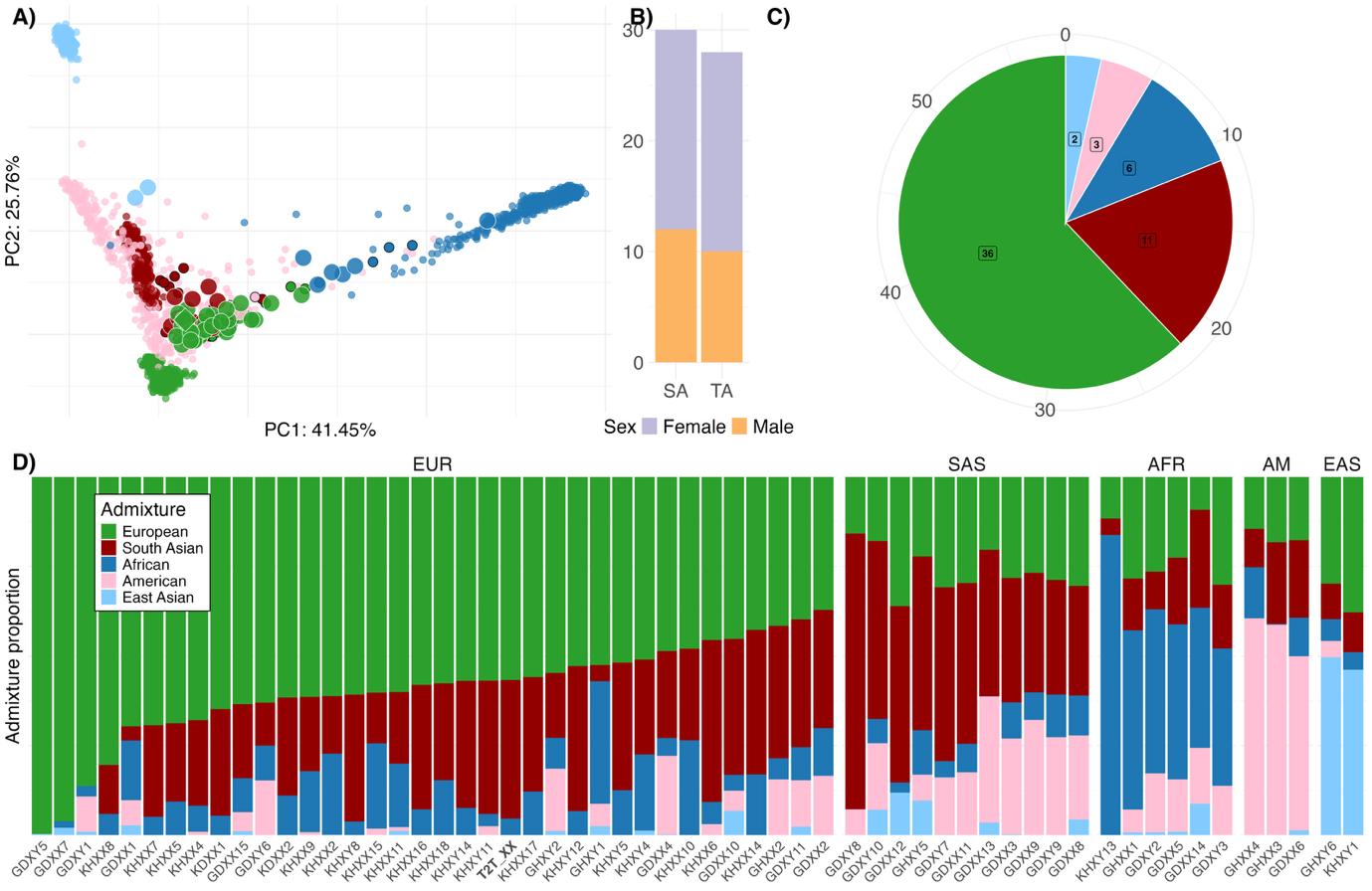

Figure 1 Basic and ancestry-based characteristics of the 58 genomes included in the Emirati genome reference. (a) The genomes are projected onto the genotype principal components computed by the 1000 Genomes phase 3 samples. Each Emirati genome is colored according to its 1000 Genomes-based global ancestry component with the maximum value. Larger dots with white borders highlight the assemblies, while in the case of trio assemblies, the parents are shown with smaller black-bordered dots. The T2T-ERG_XX sample is denoted by diamond-shaped symbols, with the T2T-ERG_XX genome located between the two parent genomes. (b) The number of pangenome single (SA) and trio (TA) assemblies stratified by sex, including the T2T assembly. (c) The distribution of the main ancestry components of the genomes across the cohort. (d) For each Emirati genome, the contributing ancestry components after supervised admixture analysis based on 1000 Genomes global populations, i.e., European, South Asian, African, American, and East Asian, c.f. Supplementary Table 17.

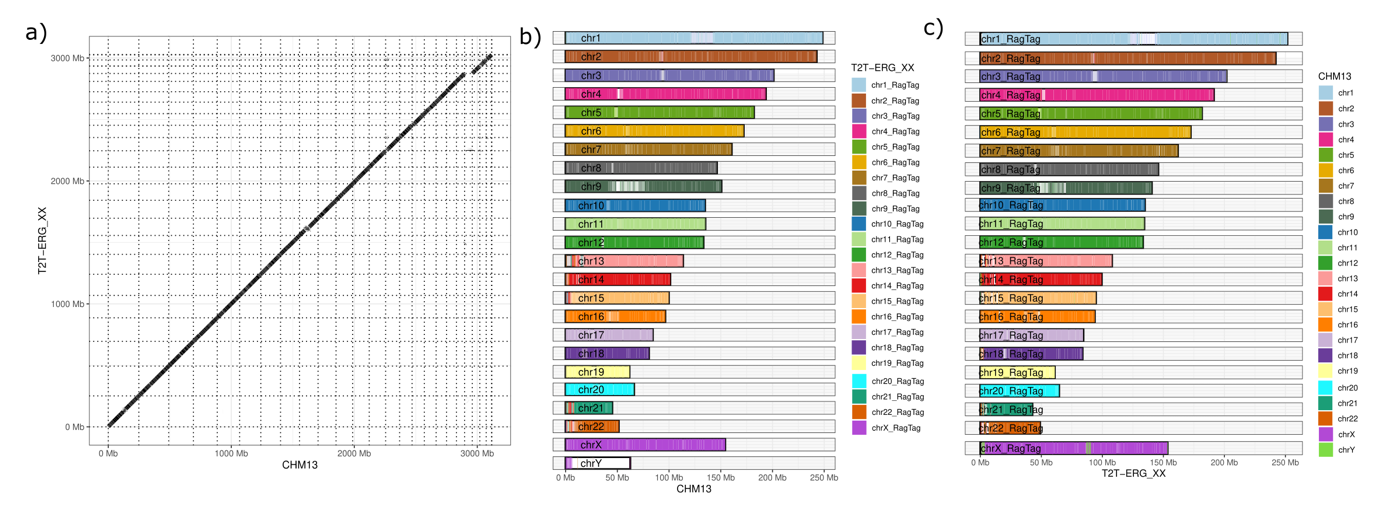

Figure 2 Alignment of T2T-ERG_XX maternal haplotype with CHM13v2.0. All primary alignments are shown. (a) Dotplot. (b) The coverage of T2T-ERG_XX chromosomes with CHM13 chromosomes. (c) The coverage of CHM13 chromosomes with T2T-ERG_XX chromosomes. In (b) and (c), all assembly sequences with a length of more than 5 Mb are shown.

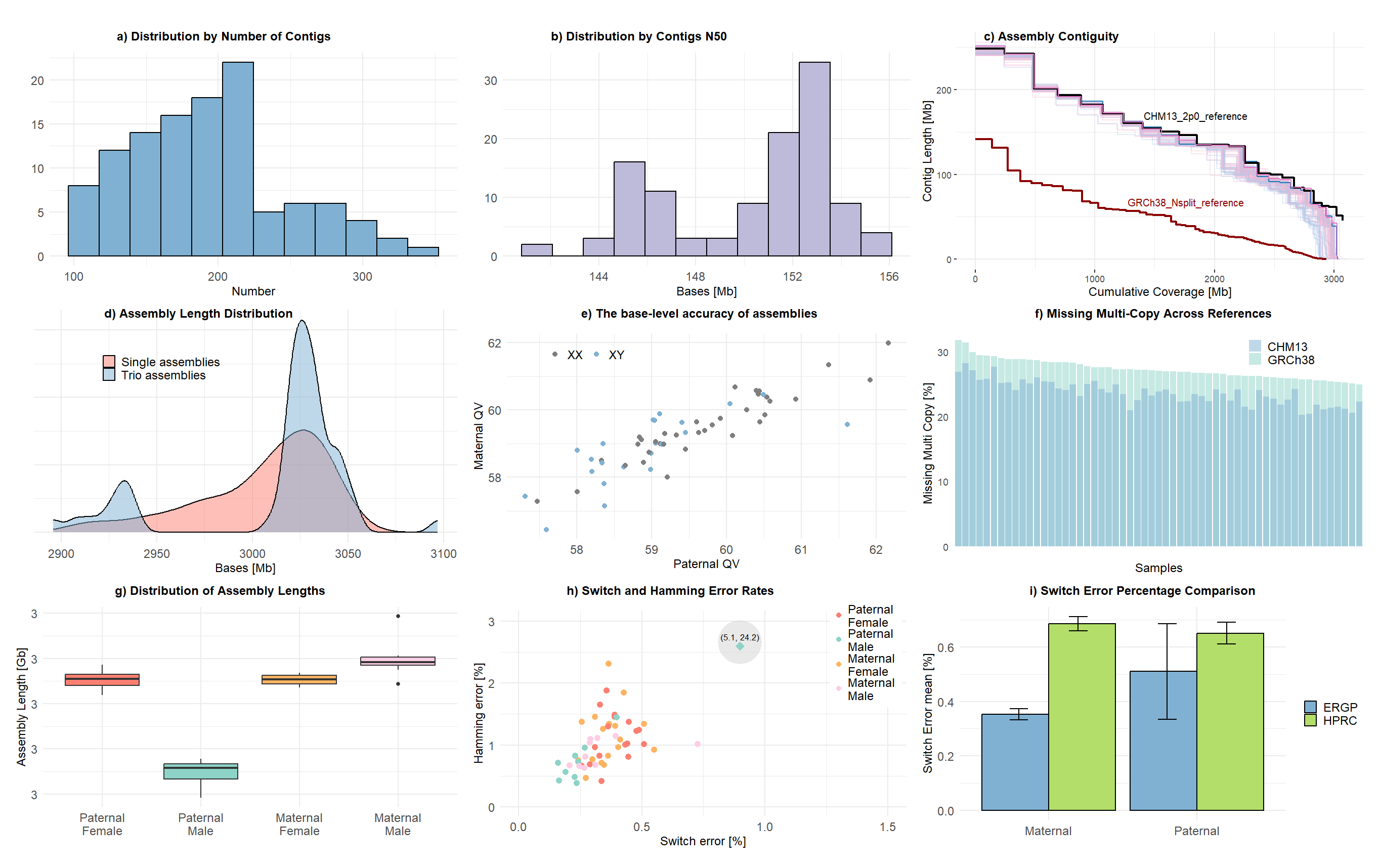

Figure 3 Assembly characteristics of 57 post-processed HiFi assemblies. Panel (a) presents the distribution by the number of contigs for all assemblies. Panel (b) shows the N50 length across assemblies. Panel (c) shows the contig length vs. cumulative assembly coverage, compared with CHM13 and GRCh38 references. Panel (d) displays the overall sequence length for 114 haplotypes, stratified by single-sample and trio-assembly approaches. Panel (e) compares the quality values (QV) between paternal and maternal haplotypes. Panel (f) illustrates the percentage of missing multi-copy genes. Panel (g) summarizes total contig lengths stratified by sex and haplotype for trio-assembly samples. Panel (h) plots Hamming error versus switch error (%) for the 27 trio assemblies. Note: the highlighted outlier corresponds to a male paternal haplotype with elevated Hamming error; see the main text for details. Panel (i) compares the mean switch error of all haplotypes from the 27 trio assemblies with the mean switch error reported by the Human Pangenome Reference Consortium (HPRC).

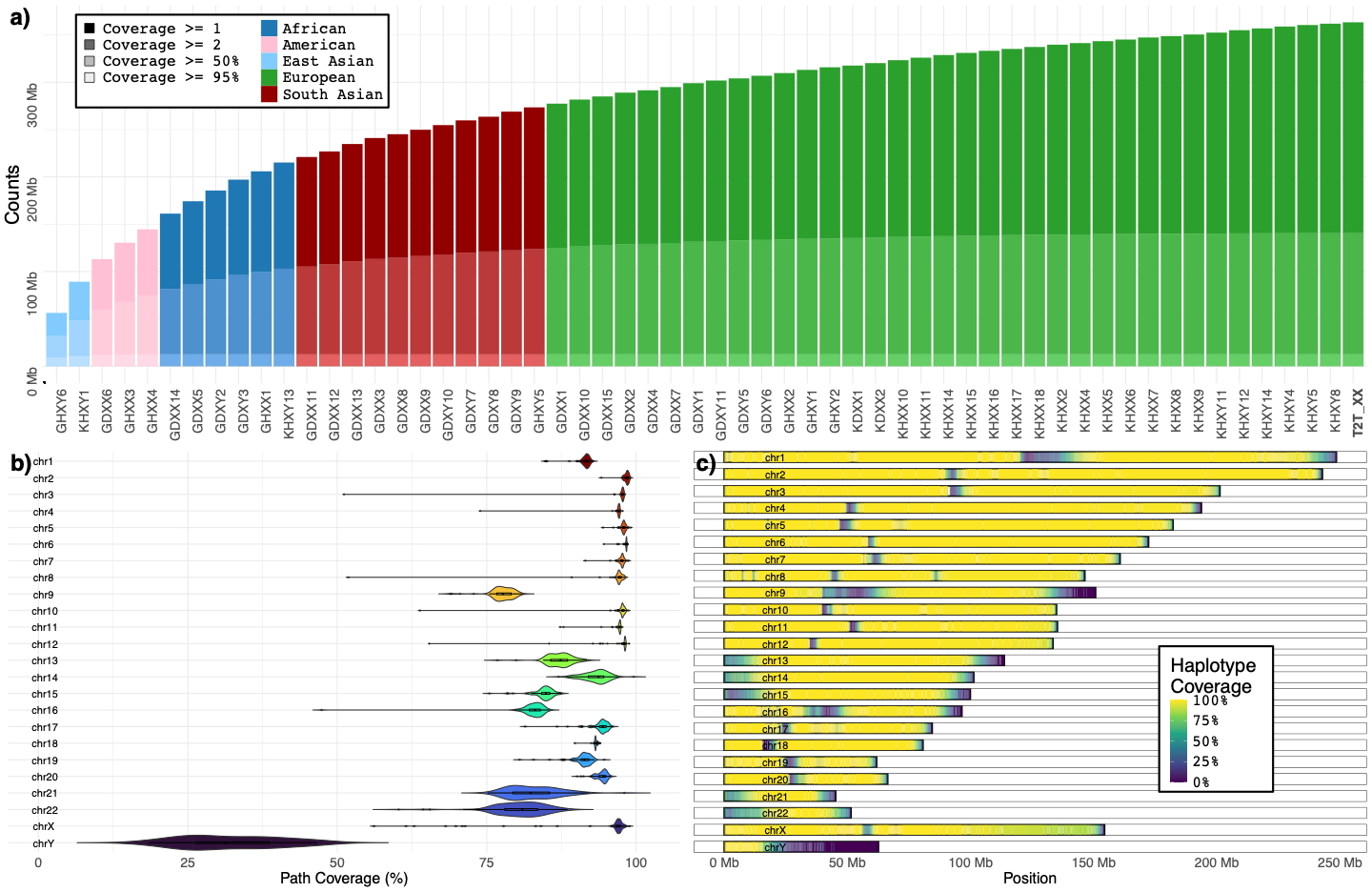

Figure 4 Characteristics of the pangenome graph (a) Bar plots depict the ordered growth of the pangenome graph in base pairs across successive sample additions. Values are reported for individual sample contributions as well as cumulative growth when samples are added in Panacus order, from 1 to 58. The growth curve demonstrates a steep initial increase in graph size with the first samples, followed by a gradual plateau, indicating diminishing contribution to the total graph as additional genomes are incorporated. (b) Violin plot of graph segment coverage across chromosomes in 116 scaffold assemblies mapped to the CHM13 reference. The median segment coverage for all autosomes ranges from 77.7% (chromosome 9) to 98.5% (chromosome 2). Coverage exceeded 97% for most large autosomes (chromosomes 2–8, 10–12), while lower median values were observed for chromosomes 9 (77.7%), 22 (80.9%), 16 (83.1%), 21 (82.3%), and 15 (84.9%). Median path coverage across sex chromosomes. The median coverage was 97.0% for chromosome X and 31.4% for chromosome Y. (c) A heatmap of path coverage across the backbone of CHM13 chromosomes, where each position is colored by the fraction of assembled haplotype contigs that traverse the graph at that locus. The pattern recapitulates the distribution summarized in panel (b), with high coverage along most autosomes and chromosome X, and pronounced depletion in repetitive regions. Coverage is lowest on the chromosome Y, consistent with its median of 31.4%, and remains high on the chromosome X, with a median of 97.0%. Reduced signal is concentrated in centromeric and subtelomeric intervals and on the short arms of the acrocentric chromosomes 13, 14, 15, 21, and 22, reflecting expected challenges in assembling highly repetitive sequences.

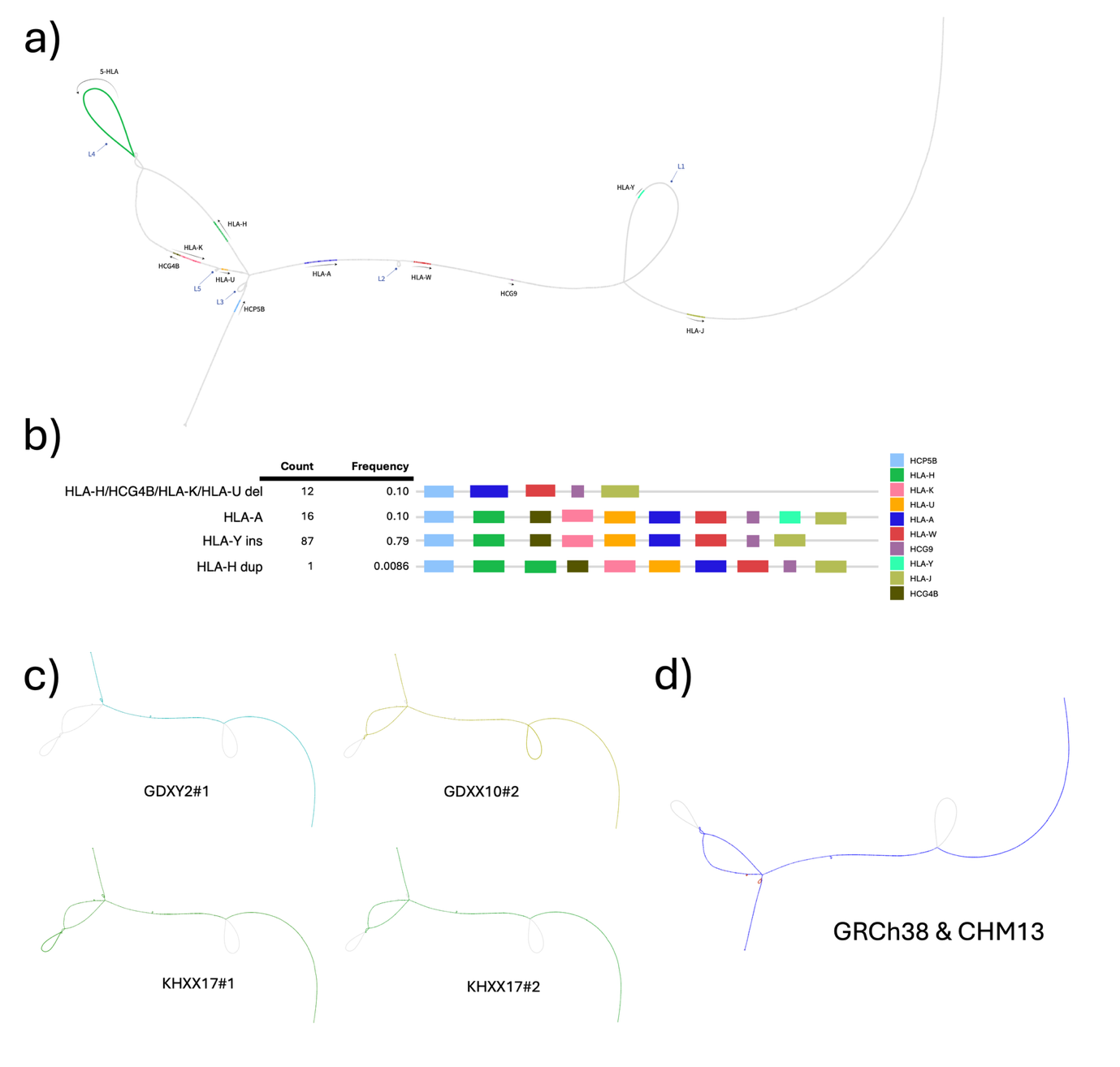

Figure 5: Visualizing complex pangenome loci. (a) Structural variation of HLA-A from the MC graph. b) Haplotype frequencies across the analyzed assemblies c-d) Representative haplotypes illustrate distinct structural configurations relative to GRCh38 & CHM13v2, including HLA-A, HLA-Y insertion, HLA-H/HCG4B/HLA-K/HLA-U deletion, and HLA-H duplication.
