## Supplementary Figures and Descriptions for "The Emirati T2T-Level Pangenome: A Graph of 58 Complete Genomes"

**Supplementary Tables**

*Supplementary Table 1:* Summary of Illumina short-read sequencing data generated using the NovaSeq 6000 platform. Sequencing was performed with a read length of 150 bp, yielding coverage exceeding 90X for all individuals. Coverage was calculated based on the CHM13 genome length.

*Supplementary Table 2:* Summary of long-read sequencing data generated for ERG_FATHER, ERG_MOTHER, and T2T-ERG_XX using PacBio HiFi and Oxford Nanopore Technologies (ONT) platforms. Coverage is computed for the genome length of CHM13. ERG_FATHER has two HiFi and one ONT run, ERG_MOTHER has three HiFi and one ONT and T2T-ERG_XX has four HiFi and 19 ONT runs. HiFi sequencing data demonstrated high coverage across all individuals, with coverage ranging from 29.52x to 34.54x. ONT sequencing data included multiple files per sample, with varying sequencing depths and coverage ranging from 0.05x to 31.83x.

*Supplementary Table 3:* Summary of processed and merged sequencing data for T2T-ERG_XX, ERG_FATHER, and ERG_MOTHER across Illumina, Oxford Nanopore Technologies (ONT), and PacBio HiFi platforms. The coverage is computed for the genome length of CHM13. Illumina data has been downsampled by 50% for each sequencing file in Supplementary Table 1 and achieved a coverage of 46.73×. ONT and HiFi data are combined from long-read sequencing data listed in Supplementary Table 2. ONT data for T2T-ERG_XX amounts to coverage of 111.81x with long-read lengths, as shown by a maximum read length of 824,775 bp and an N50 of 16,928 bp. HiFi sequencing provided high-accuracy data with coverage values ranging from 63.01x to 126.64x and N50 values between 15,167 bp and 17,934 bp.

*Supplementary Table 4:* Summary of assembly strategies and performance metrics for maternal and paternal haplotypes of T2T-ERG_XX calculated before manually fixing one assembly error in the final maternal and paternal haplotype. The strategy incorporated combinations of assembly tools, including Verkko, ntLink (HiFi and ONT), RagTag, hifiasm, and quarTeT, with and without Ratatosk correction. The number of contigs ranged from 25 to 195, with total base counts exceeding 3.02 Gb for all assemblies. Maternal haplotypes achieved N50 values up to 153.4 Mb and QV scores over 60, while paternal haplotypes exhibited N50 values up to 162.2 Mb. Hamming error rates ranged from 7.2962 to 18.1856, highlighting variability across strategies.

*Supplementary Table 5:* Analysis of maternal and paternal T2T-ERG_XX haplotypes derived from a single individual's genome using combined Verkko and HiFiASM assemblies. Metrics from QUAST, yak, and Inspector (see Methods) include contiguity, assembly size, alignment quality, and structural correctness. The maternal haplotype contains 80 contigs (largest: 251.6 Mb, N50: 153.4 Mb), while the paternal haplotype has 100 contigs (largest: 247.6 Mb, N50: 162.2 Mb). Both haplotypes achieved high genome coverage (94.38% maternal, 94.40% paternal) and minimal duplication rates (1.017–1.018). Gene coverage, based on CHM13v2.0 annotation, and alignment metrics (NA50, NGA50) indicate high completeness and alignment quality.

*Supplementary Table 6:* Analysis of maternal and paternal T2T-ERG_XX haplotypes, detailing the start and end positions of gaps across chromosomes and their respective gap sizes. The table provides information on gaps identified in both haplotypes, spanning autosomal chromosomes, sex chromosomes, and mitochondrial DNA. Gap lengths range from 1 bp (e.g., T2TXXmatChrM) to 100,000 bp (e.g., chr19_RagTag for the paternal haplotype). Maternal gaps are predominantly small, with most ranging between 100 bp and 7,500 bp, and are distributed across chromosomes such as chr1_RagTag, chr2_RagTag, and chrX_RagTag. In contrast, paternal haplotypes show a similar distribution but include a larger gap, with the 100,000 bp gap on chr19_RagTag.

*Supplementary Table 7:* Sequencing data quality metrics for the HiFi data underlying pangenome assemblies across 27 trio (TA) and 30 single (SA) sample types for male (XY) and female (XX) individuals. Coverage values range from 25.53× to 52.42×, with ten samples marked for sequencing top-up to reach the 25x coverage cutoff.

*Supplementary Table 8:* The raw assembly characteristics of 114 haplotypes from 57 samples were categorized by sample ID, sex, and type (TA: trio assembly or SA: single assembly). Metrics were generated using QUAST, Yak, and Inspector, as described in the methods. The table includes contiguity metrics such as the number of contigs per haplotype, which range from 233 to 836, and N50 values spanning 25.5 Mb to 85.3 Mb. Total genome lengths (≥0 bp) are consistent across assemblies, ranging from 2.89 Gb to 3.09 Gb, with a uniform GC content reported between 40.73% and 40.88%. The largest contigs range in size from approximately 89.6 Mb to 199.0 Mb, with a median of 131.4 Mb. Mismatches per 100 kbp are low, ranging from 125.9 to 177.4 per 100 kbp. Genome alignment metrics are high, with genome fractions exceeding 90.18% and the largest alignments spanning 138.4 Mb. Duplication ratios are stable at around 1.014–1.033, and structural error metrics, such as switch error rates and Hamming error rates, remain below 1%.

*Supplementary Table 9:* The assembly characteristics of 114 haplotypes from 57 samples were categorized by sample ID, sex, and type (TA: trio assembly or SA: single assembly). Metrics were generated using QUAST, Yak, and Inspector after scaffolding, polishing, and error correction. The table includes contiguity metrics such as the number of contigs per haplotype, which range from 100 to 335, and N50 values spanning 140.9 Mb to 154.9 Mb. Total genome lengths (≥0 bp) are consistent across assemblies, ranging from 2.89 Gb to 3.09 Gb, with a uniform GC content reported between 40.73% and 40.88%. The largest contigs range in size from approximately 240.3 Mb to 255.5 Mb, with a median of 243.2 Mb. Mismatches per 100 kbp are low, ranging from 124.0 to 177.2 per 100 kbp. Genome alignment metrics are high, with genome fractions exceeding 90.22% and the largest alignments spanning 147.7 Mb. Duplication ratios are stable at around 1.012–1.031, and median structural error metrics, such as switch error rates and Hamming error rates, remain below 1%.

*Supplementary Table 10:* Comparison of graph assembly metrics across chromosomes for a cohort of 58 samples, with different backbones (GRCh38 and CHM13v2). The table details the number of edges, nodes, and graph lengths for each dataset. Edges range from 2,270 (chrM, CHM13v2 backbone) to 9.2 million (chr1, CHM13v2 backbone), with the total graph containing 121 million edges for CHM13v2 backbone, 111 million for GRCh38 backbone. Nodes range from 1,687 (chrM, CHM13v2 backbone) to 6.7 million (chr1, CHM13v2 backbone), with total node counts of 88 million (CHM13v2 backbone) and 79 million (GRCh38 backbone). Graph lengths span from 17,312 bp (chrM, CHM13v2 backbone) to 2667.4 million bp (chr1, CHM13v2 backbone), with total graph lengths of 3.35 Gb (CHM13v2 backbone), and 3.23 Gb (GRCh38 backbone).

*Supplementary Table 11:* Pangenome graph growth curve data underlying Figure 4a illustrates the ordered growth of the graph in base pairs (bp) for multiple samples. Metrics are shown for single-sample growth, incremental growth with additional samples (e.g., 1, 2, 3, up to 58 genomes). The data includes growth curves for panacus-based ordered-growth increments. Growth metrics reveal initial rapid expansions with fewer samples and a plateau as additional samples contribute less to the graph's total base pairs.

*Supplementary Table 12:* The number of variants in the pangenome graph incorporating the maternal Emirati T2T haplotype (T2T-ERG_XX) was calculated for 57 Emirati genomes. The table reports the total number of variant records, categorized into SNPs, MNPs, indels, other variant types, multiallelic sites, and multiallelic SNP sites. Comparisons between graphs with different backbone reference highlight the impact of integrating these references on the graph. Variants specified with respect to CHM13 backbone consistently show the highest numbers across categories, reflecting its completeness.

*Supplementary Table 13:* Minigraph-Cactus graph paths for the Emirati T2T pangenome graph with CHM13 backbone.

*Supplementary Table 14:* Comparison of CHM13 and T2T-ERG_XX T2T chromosome assembly metrics for maternal (ERG_XX#2) and paternal (ERG_XX#1) T2T haplotypes. The table includes the mapping between CHM13 chromosomes and corresponding T2T chromosomes (RagTag paths), along with path length, percentage of CHM13 coverage, T2T and CHM13 chromosome lengths, and the number of gaps in each T2T chromosome. Path lengths vary from approximately 35 Mb (chr21) to 239 Mb (chr2), with longer path lengths aligning to larger chromosomes such as chr1 and chr2. CHM13 coverage ranges from 77.78% (chr21, ERG_XX#2) to 98.86% (chrX, ERG_XX#2), with most chromosomes exhibiting high coverage (>90%), indicating strong alignment between CHM13 and T2T assemblies. T2T chromosome lengths range from 38 Mb (chr21) to 251 Mb (chr1), closely matching CHM13 chromosome lengths, which span from 45 Mb (chr21) to 248 Mb (chr1). The number of gaps within T2T chromosomes ranges from 0 to 13, with most chromosomes containing fewer than 5 gaps, demonstrating high assembly continuity overall.

*Supplementary Table 15:* Longest contig path per pangenome haplotype in relation to CHM13 chromosomes across all 58 assemblies. The percentage indicates the proportion of each contig assigned to its respective CHM13 chromosome relative to the total chromosome length. Autosomes show a median coverage of 94.55% (range: 46.38% to 99.33%). Chromosome X, in female haplotypes and male haplotype 2, exhibits a median coverage of 96.98% (range: 27.63% to 98.82%). Chromosome Y in male haplotype 1 has a lower median coverage of 30.71% (range: 14.42% to 46.57%), indicating persistent fragmentation in the current assemblies.

*Supplementary Table 15:* SNP counts for 58 pangenome assemblies as reported by the Minigraph-Cactus pipeline for the CHM13 backbone and GRCh38 backbone graph with respect to CHM13 and GRCh38 are provided. For comparison, the number of SNP calls obtained by HiFi sequencing data-based SNP calling with Sention with respect to CHM13 and GRCh38 is provided. Ordering is according to the growth curve in Figure 3, i.e., the main ancestry component, and then lexicographically.

*Supplementary Table 16:* Overview of sample metadata, including sex, assembly type (single assembly (SA) or trio assembly (TA)), ancestry proportions (European (EUR), South Asian (SAS), American (AMR), African (AFR), and East Asian (EAS)), and the main global ancestry component for each genome, defined as the ancestry with the highest proportional value. The table summarizes data for 57 genomes from two sources (G42 and KU). European ancestry is the most prevalent ancestral component across the genomes. The ancestry proportions vary widely, with European contributions reaching as high as 99.7% in GDXY5 and African contributions as high as 83.8% in KHXY13.

*Supplementary Table 17:* HLA haplotypes stratified by maternal and paternal assembly haplotype. The table summarizes the occurrence of major structural haplotypes, including the HLA-Y insertion, the multi-gene deletion spanning HLA-H, HCG4B, HLA-K, and HLA-U, and the rare HLA-H insertion haplotype. Counts and frequencies are reported separately for maternal and paternal haplotypes.

*Supplementary Table 18:* Assessment of different linear and graph-based genomic references using Illumina short-read whole-genome sequencing data from n=119 Emirati individuals generated using the NovaSeq 6000 platform. The table evaluates mapping, coverage, duplication rates, and variant calling metrics across multiple references, including GRCh38, CHM13, and graph-based pangenomes. These graphs are the EPRC graphs (with GRCh38 and CHM13 backbones) and the HPRC graph with a GRCh38 backbone. Coverage was calculated using the CHM13 genome size, which ranges from 30.79X to 76.84X with a median of 40.00X. Mapping rates were high across all references, with median mapped read percentages ranging from 91.26% (EGR graph GRCh38) to 92.54% (CHM13), and duplication rates varied by reference. Median mapped base percentages ranged from 89.96% (EGR graph GRCh38) to 91.79% (CHM13), indicating consistent alignment performance. The variant analysis identified mean counts of 4.06 million SNVs and 967,104 indels across references.

*Supplementary Table 19:* Performance metrics for variant calling using short- and long-read NGS data across three references (linear GRCh38, KU pangenome, and HPRC pangenome) against GIAB truth sets (HG001–HG007). Evaluation metrics include recall, precision, F1 Score, and Matthews Correlation Coefficient (MCC), along with detailed counts for true positives (TP), false negatives (FN), false positives (FP), and unknowns (UNK). Additional statistics include TiTv ratios, heterozygous-to-homozygous (het/hom) ratios, the fraction of variants not assessed (Frac NA), and BED-based true negatives (TN). Overall, long-read data generally yield higher precision and recall for indels, while SNV calling across all references shows high performance (F1 > 0.99). Among references, the linear GRCh38 tends to achieve slightly higher MCC and lower error rates for SNVs, while the pangenome-based references provide competitive or better results for indel detection in some cases. These findings highlight both the advantages of long-read sequencing and the subtle performance differences between reference choices.

*Supplementary Table 20:* List of computational tools, software packages, and their corresponding versions used in the respective analyses throughout this study. The table details the specific role of each tool (e.g., assembly, alignment, variant calling, graph construction, phasing, and visualization), along with version numbers and references, to ensure reproducibility and transparency of all analyses performed.

**Supplementary Figures**

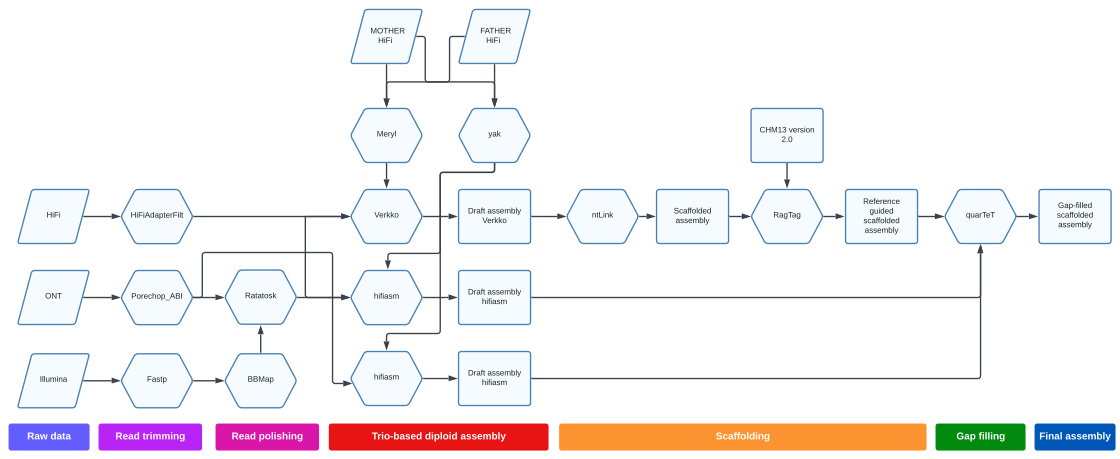

Supplementary Figure 1: Processing workflow for human chromosome-level genome assemblies. The diagram outlines steps for integrating sequencing data from multiple platforms (HiFi, ONT, and Illumina) to create a comprehensive trio-based diploid assembly. Following read trimming (purple) and long-read polishing (pink), the assembly stage (red) begins by generating k-mer databases of parental reads using Meryl and Yak for the assembly pipelines Verkko and hifiasm, respectively. The generated draft assemblies are scaffolded (orange) using ntLink and further improved using RagTag for reference-guided assembly based on the CHM13v2.0 reference genome. The final assembly refinement involves gap-filling (green) using quarteT, culminating in the gap-filled scaffolded assembly (dark blue).

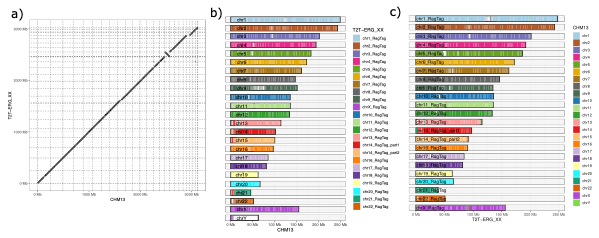

Supplementary Figure 2: Alignment of T2T-ERG_XX paternal haplotype with CHM13v2.0. All primary alignments are shown. (a) Dotplot. (b) The coverage of T2T-ERG_XX chromosomes with CHM13 chromosomes. (c) The coverage of CHM13 chromosomes with T2T-ERG_XX chromosomes. In (b) and (c), all assembly sequences with a length of more than 5 Mb are shown.

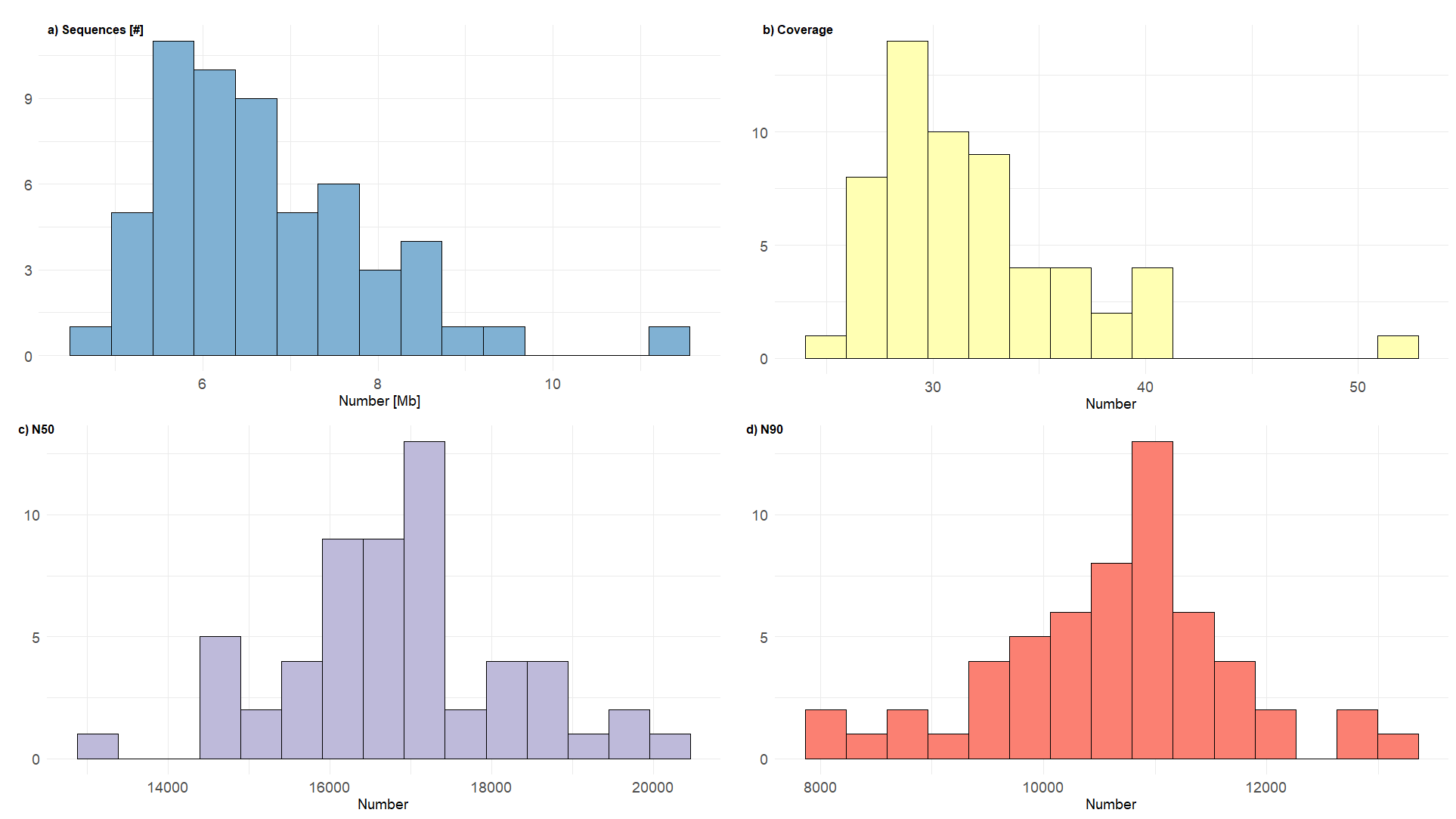

Supplementary Figure 3: Sequencing summary quality control across the HiFi data of the 57 Emirati genome reference genomes. The figure summarizes quality control (QC) results from FastQC analysis showcasing distributions across multiple QC metrics. (a) Sequence number; (b) Average genome coverage; (c) Read N50 length; (d) Read N90 length.

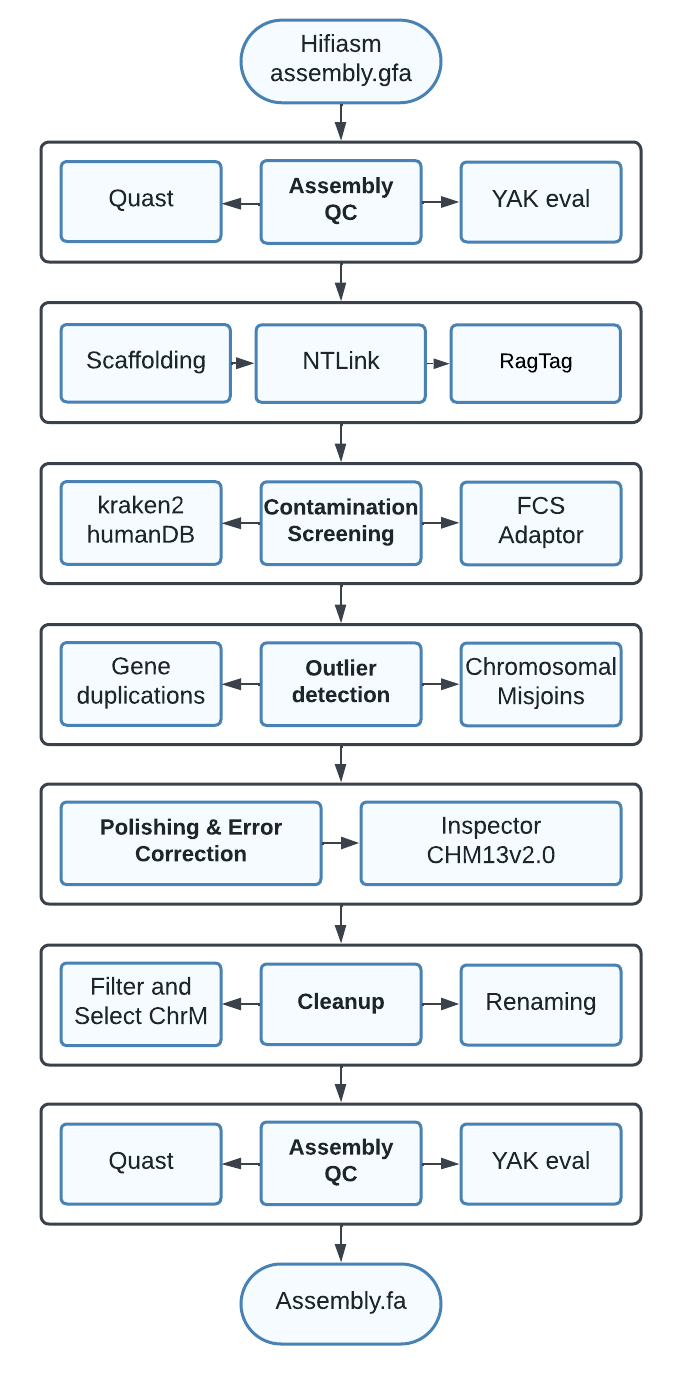

*Supplementary Figure 4: Workflow for post-assembly processing of hifiasm-generated assemblies in pangenome construction. The initial assembly undergoes quality control (QC) using Quast and YAK evaluation. Scaffolding is performed with ntLink and further improved using RagTag for reference-guided assembly based on the CHM13v2.0 reference genome, followed by contamination screening using Kraken2 with the human database and FCS Adaptor to identify contaminants. Outlier detection assesses chromosomal misjoins and gene duplications. Polishing and error correction are executed with reference to CHM13v2.0 using Inspector. Cleanup steps include filtering and selecting mitochondrial chromosomes (chrM) and renaming. The final assembly is subjected to another round of QC with Quast and YAK evaluation to ensure quality and accuracy. of QC with Quast and YAK evaluation to ensure quality and accuracy.*

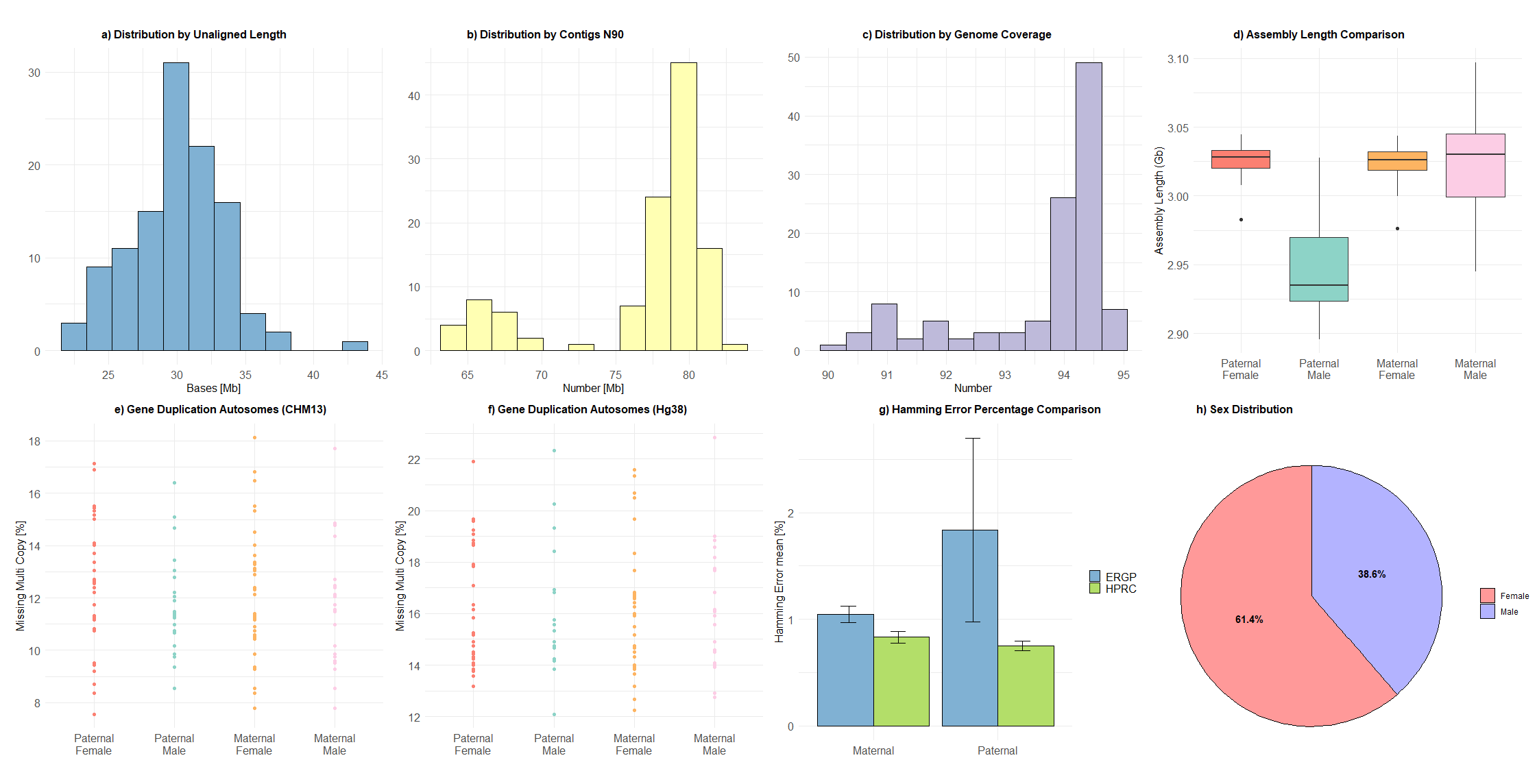

Supplementary Figure 5: The figure shows evaluations of genome assembly and quality assessment across various panels. Panel (a) presents the distribution of unaligned contig lengths in megabases (Mb). Panel (b) depicts the N90 statistic for contig sizes (Mb). Panel (c) shows the histogram of genome coverage for all 114 haplotypes. Panel (d) compares the assembly length (Gb) across maternal and paternal haplotypes for single-sample and trio-assembly strategies. Panel (e) and Panel (f) display percentages of missing multi-copy regions with respect to GRCh38 and CHM13, respectively. Panel (g) contrasts Hamming error rates (mean percentage) between maternal and paternal genomes for trio samples, as evaluated by the Emirati T2T pangenome and HPRC datasets. Panel (h) summarizes the gender composition of the analyzed dataset in percentage values, indicating the proportions of male and female samples.

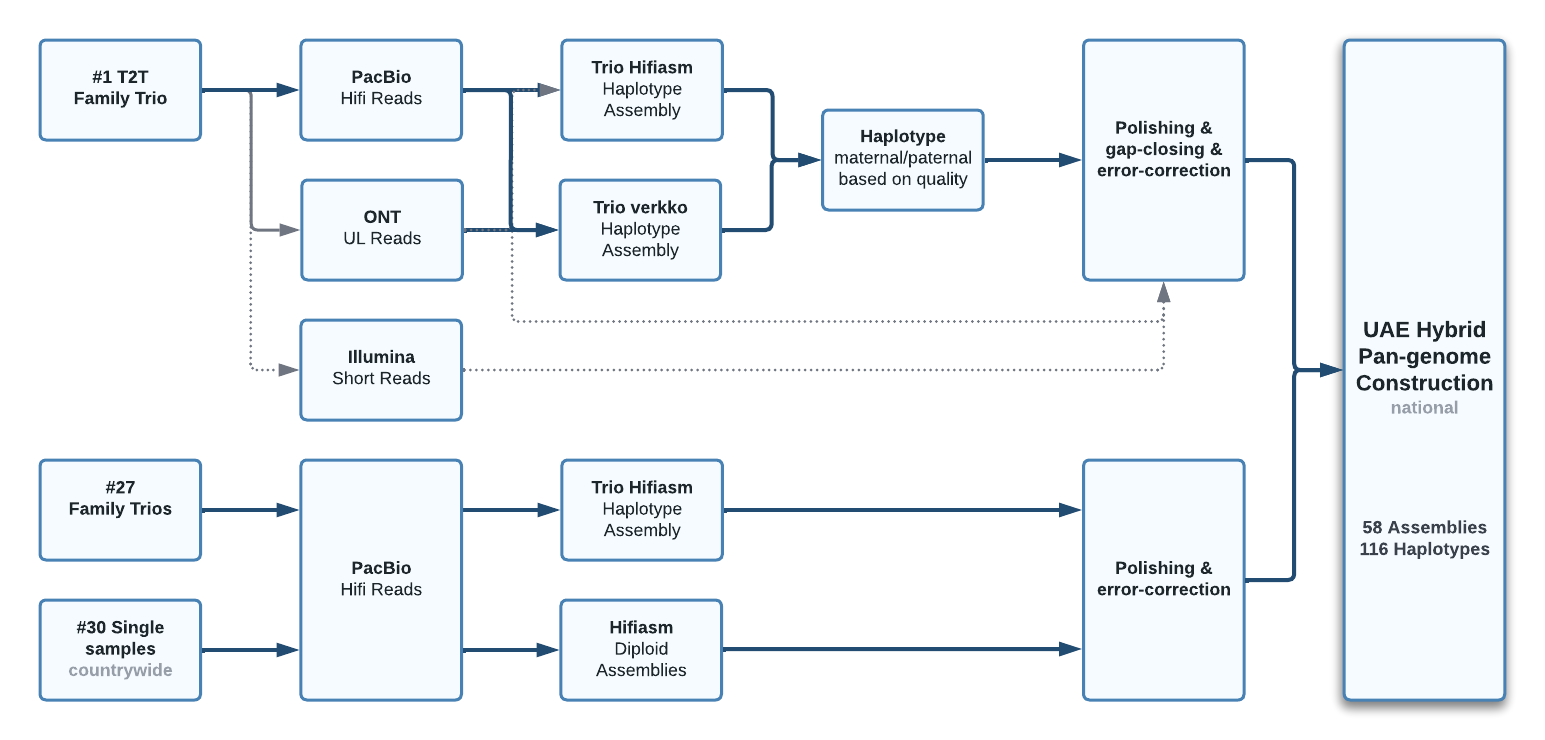

Supplementary Figure 6: Workflow for the Emirati T2T pangenome graph construction, integrating sequencing and assembly of a T2T family trio, 27 family trios, and single samples using PacBio HiFi, ONT ultra-long, and Illumina reads. Haplotype assemblies are generated with trio hifiasm and trio Verkko, while hifiasm produces diploid assemblies for single samples. Maternal and paternal haplotypes are selected based on quality, followed by polishing, gap-closing, and error correction. The final Emirati T2T pangenome comprises 58 assemblies and 116 haplotypes, capturing the genetic diversity of the UAE population.

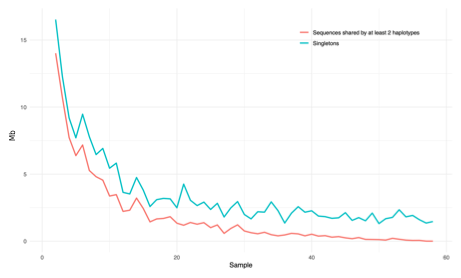

*Supplementary Figure 7: The novel sequence in Mb which is added by each of the pangenome assemblies, i.e. singleton sequence not present in the assemblies added to the graph before, as well as the sequence that is added, which has been represented by only one other assembly before. The last assembly added is T2T-ERG_XX, which ads no new sequence that had not been represented at least by two assemblies before, but still introduces 1.46 Mb singleton sequence (cf. Supplementary Table 11).*

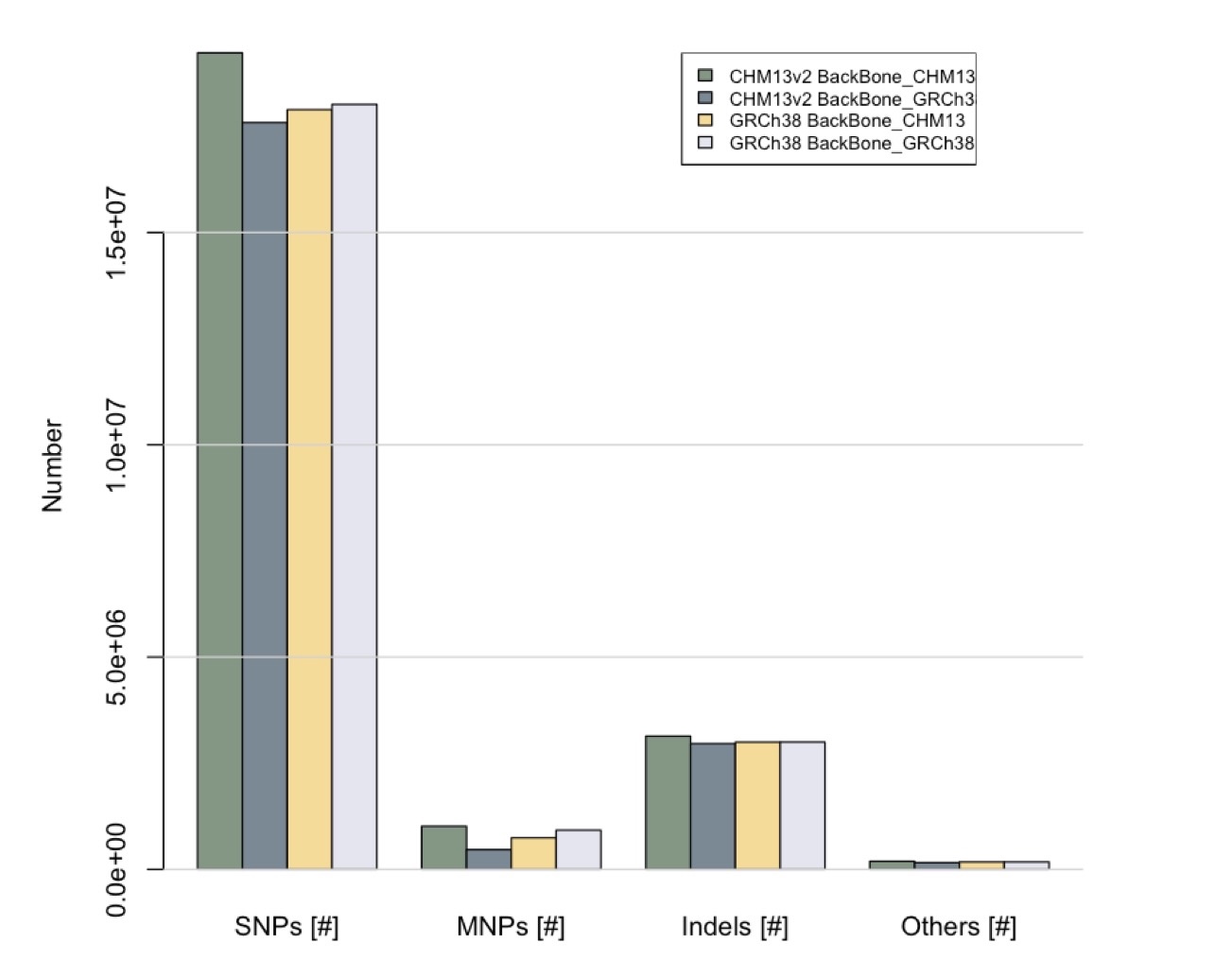

*Supplementary Figure 8: The number of variants of the Emirati pangenome assemblies deduced from the Emirati T2T pangenome graphs with backbone CHM13 and GRCh38, respectively. In both graphs, the variants were counted with respect to both CHM13 and GRCh38 and the cohort variant counts across the cohort compared. The bar plot stratifies SNPs, MNPs, indels, complex variants, multiallelic sites, and multiallelic SNP sites. Comparisons across graphs constructed with different backbone references demonstrate the influence of reference choice. Graphs using the CHM13 backbone consistently show the highest number of total variants across all categories, reflecting the improved resolution of this reference structure.*

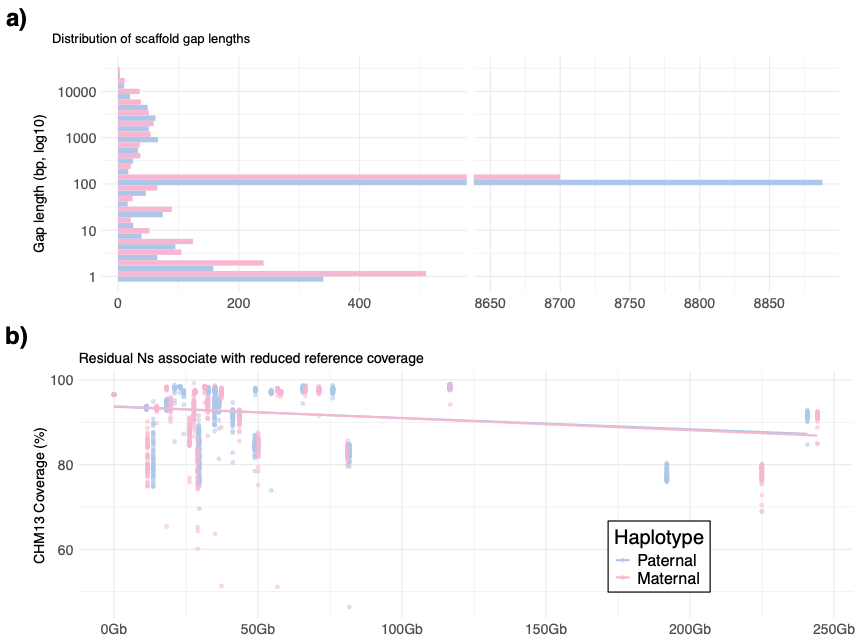

Supplementary Figure 9: N regions across haplotypes, their size distribution, and their link to reference coverage. Panel a) displays the distribution of gap lengths on a log10 scale across primary contigs, with an axis break to reveal the long tail of rare large gaps. Panel b) relates total N content per haplotype chromosome to percent coverage of the CHM13 reference path, with a linear fit and points colored by haplotype. Across 57 individuals and 114 haplotypes, 2,679 primary contigs were evaluated, of which 2,584 contained at least one gap and 95 were gap-free, representing approximately 3.37 × 10^11 bp mapped to CHM13 in total, and an aggregate gap content of about 0.00115 percent.

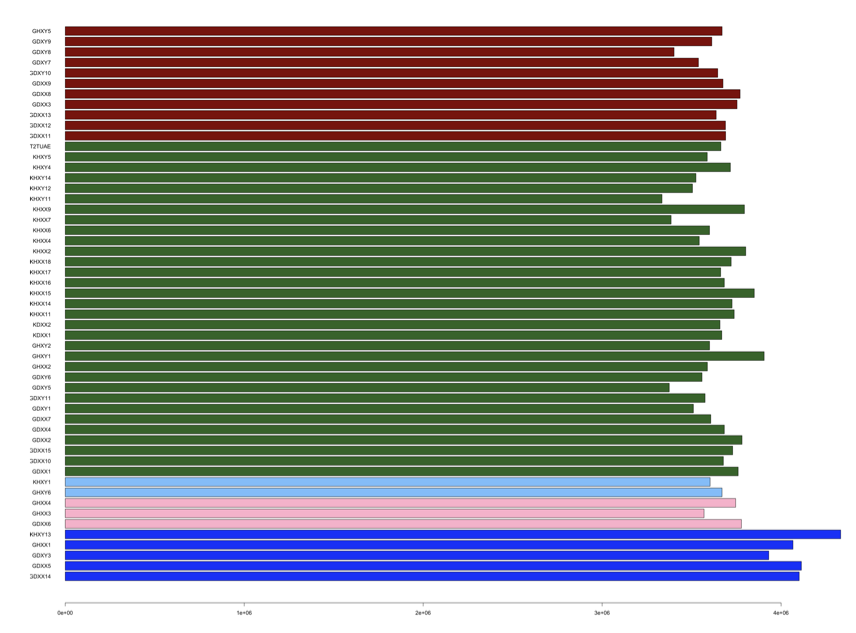

*Supplementary Figure 10: Number of SNVs per pangenome genome within the Emirati T2T pangenome graph, coloured by the genome’s main global ancestry component.*

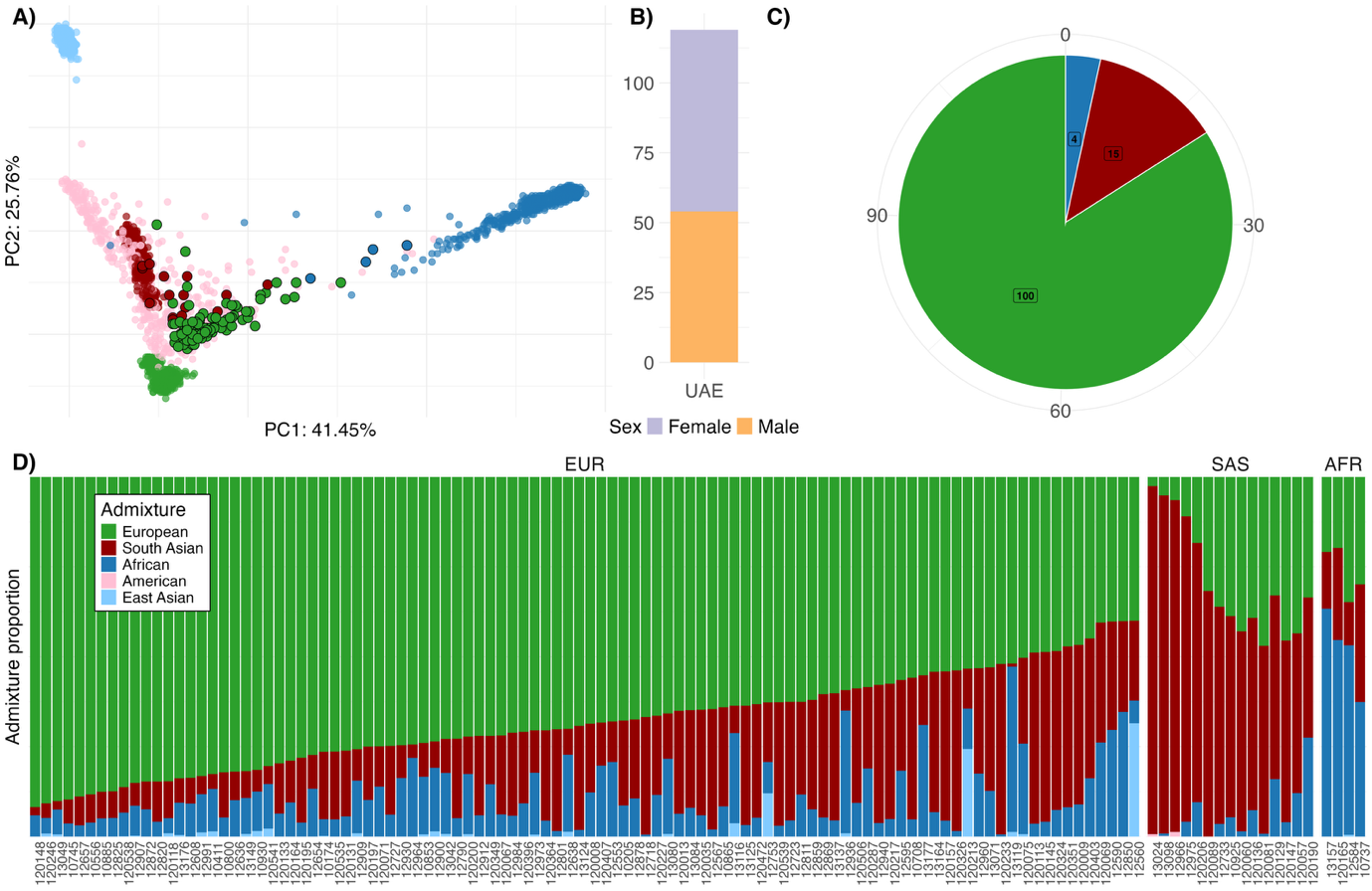

Supplementary Figure 11: Basic and ancestry-based characteristics for a cohort of 119 Emirati individuals with Illumina whole genome sequencing data used for graph evaluation. (a) The genomes are projected onto the genotype principal components computed by the 1000 Genomes phase 3 samples. Each sample is colored according to its 1000 Genomes-based global ancestry component, with the maximum value indicated. (b) Shows the cohort stratified by sex. (c) The distribution of the main ancestry components of the genomes across the cohort. (d) The contributing ancestry components for each sample, as determined by supervised admixture analysis based on the 1000 Genomes global populations, i.e., European, South Asian, African, American, and East Asian.

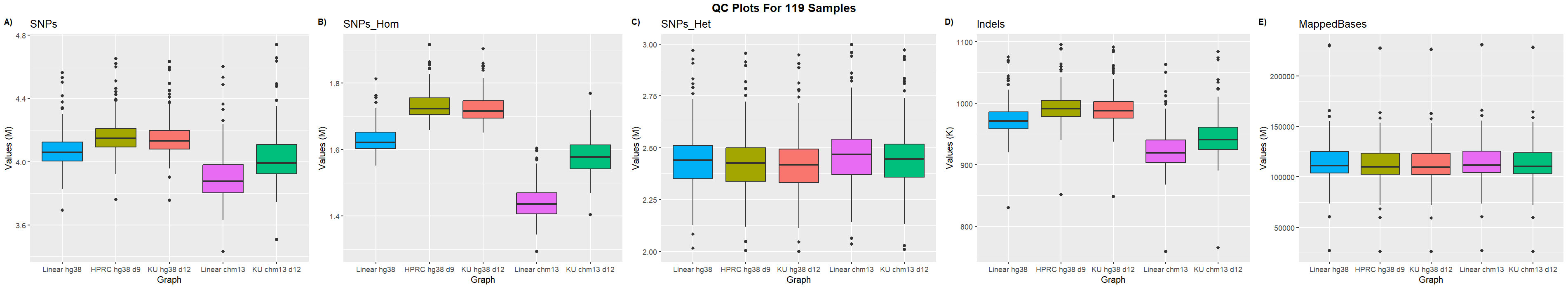

Supplementary Figure 12: Variant calling metrics for 119 UAE Illumina short-read samples aligned to linear and graph-based reference genomes, including GRCh38, CHM13v2, the public HPRC graph with GRCh38 backbone (HPRC_hg38_d9), the Emirati T2T pangenome graph with hg38 backbone (KU_hg38_d12), and the Emirati T2T pangenome graph with CHM13v2 backbone. Graph-based references generally resulted in a higher number of variant calls compared to their linear counterparts. CHM13-based references consistently produced fewer SNPs and indels than hg38-based references, with notably fewer homozygous SNPs, whereas hg38-based references produced fewer heterozygous SNPs. Among the graph-based references, the public HPRC_hg38_d9 graph called slightly more variants and showed a marginally higher number of mapped bases than the KU_hg38_d12 graph. Linear reference alignments showed a slightly higher overall number of mapped bases compared to graph-based alignments.

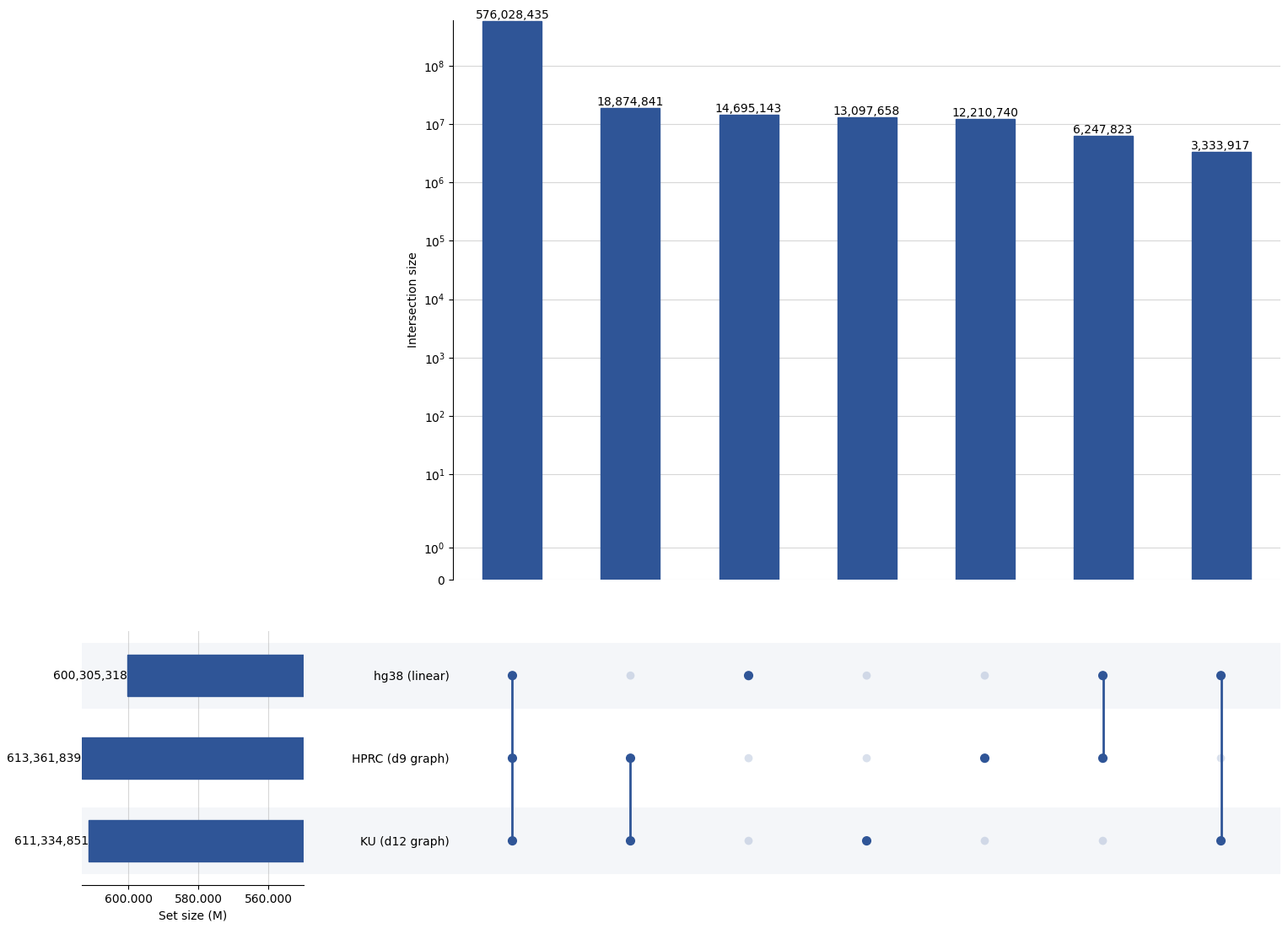

Supplementary Figure 13: Intersection analysis of variant calls from 119 UAE Illumina short-read samples aligned to linear and graph-based reference genomes, including hg38, the public HPRC graph with hg38 backbone (HPRC_hg38_d9), and the Emirati T2T pangenome graph with hg38 backbone. The UpSet plots reveal three dominant groups: (1) variants consistently identified by all references, (2) variants uniquely called by the graph-based references, and (3) variants exclusively detected by the linear reference but absent in the graphs. Across all references, a total of 644,488,557 unique variants were identified. Graph-based references called at least 11 million more variants than the linear hg38 reference across the aggregate dataset. The majority of variants, 576,028,435 (89.37%), were shared across all reference types. The linear reference exclusively contributed 2.28% of variants, while the graph-based references shared 2.92% of variants not present in the linear reference.

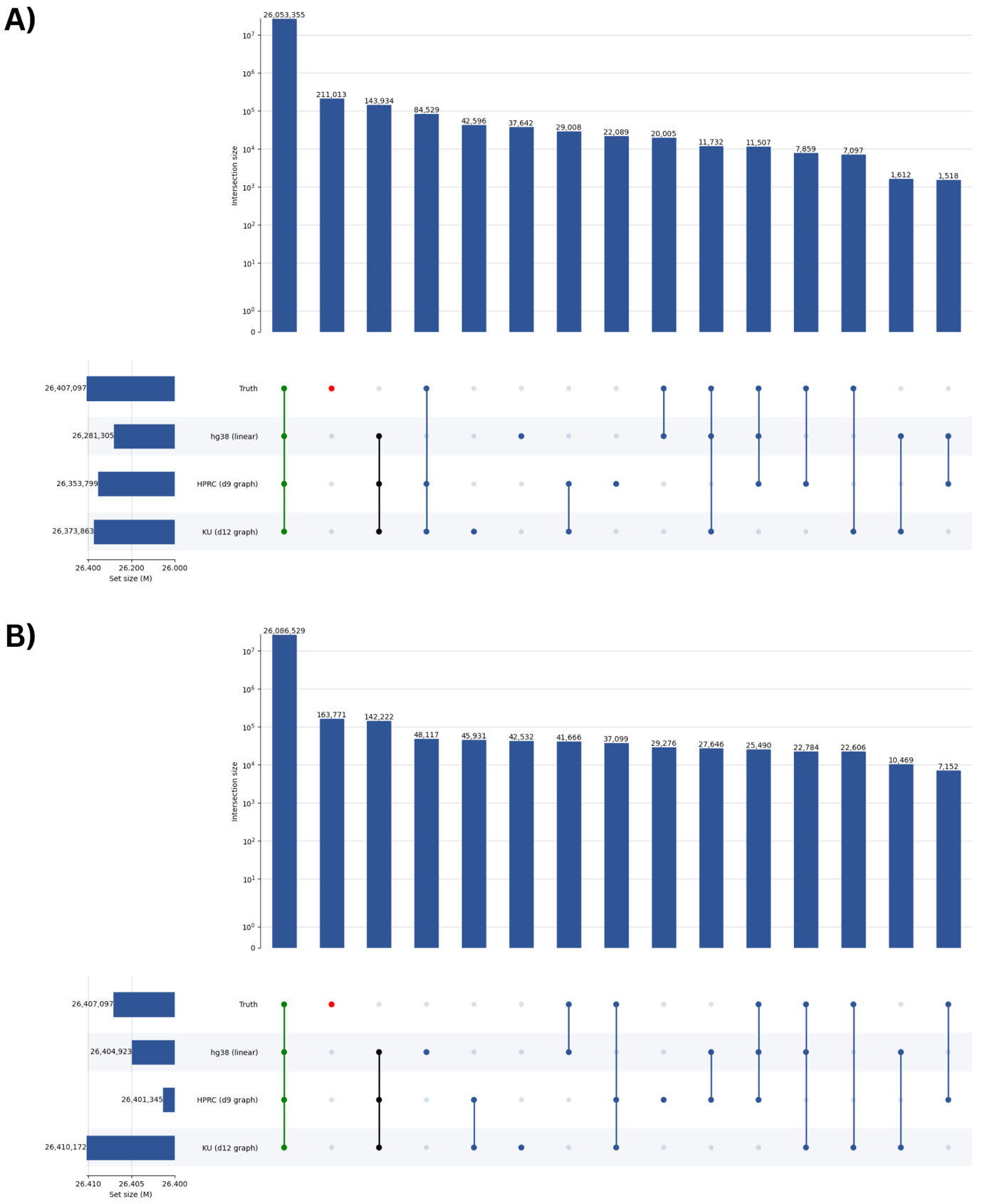

Supplementary Figure 14: Intersection analysis of variant calls from well-characterized Genome in a Bottle (GIAB) samples (HG001–HG007), aligned to linear and graph-based reference genomes, including hg38, the public HPRC graph with hg38 backbone (HPRC_hg38_d9), and the Emirati T2T pangenome graph with hg38 backbone. (A) Results based on Illumina short-read data. (B) Results based on PacBio long-read data. The GIAB truth set contains 26,407,097 variants across the seven samples. Of these, 98.66% are recovered by all references using short reads, and 98.78% using long reads. For short-read alignments, 211,013 variants from the truth set remain uncalled by any reference, compared to 163,771 in the long-read analysis. Notably, approximately 0.5% of variants consistently called across all references are not present in the truth set for both technologies. Among short-read results, the fourth largest variant group (84,529 variants, 0.32%) consists of truth-set variants called by the graph references but missed by the linear hg38 reference. In long-read data, this group drops to the eighth largest, likely reflecting improved alignment specificity with long-read inputs.

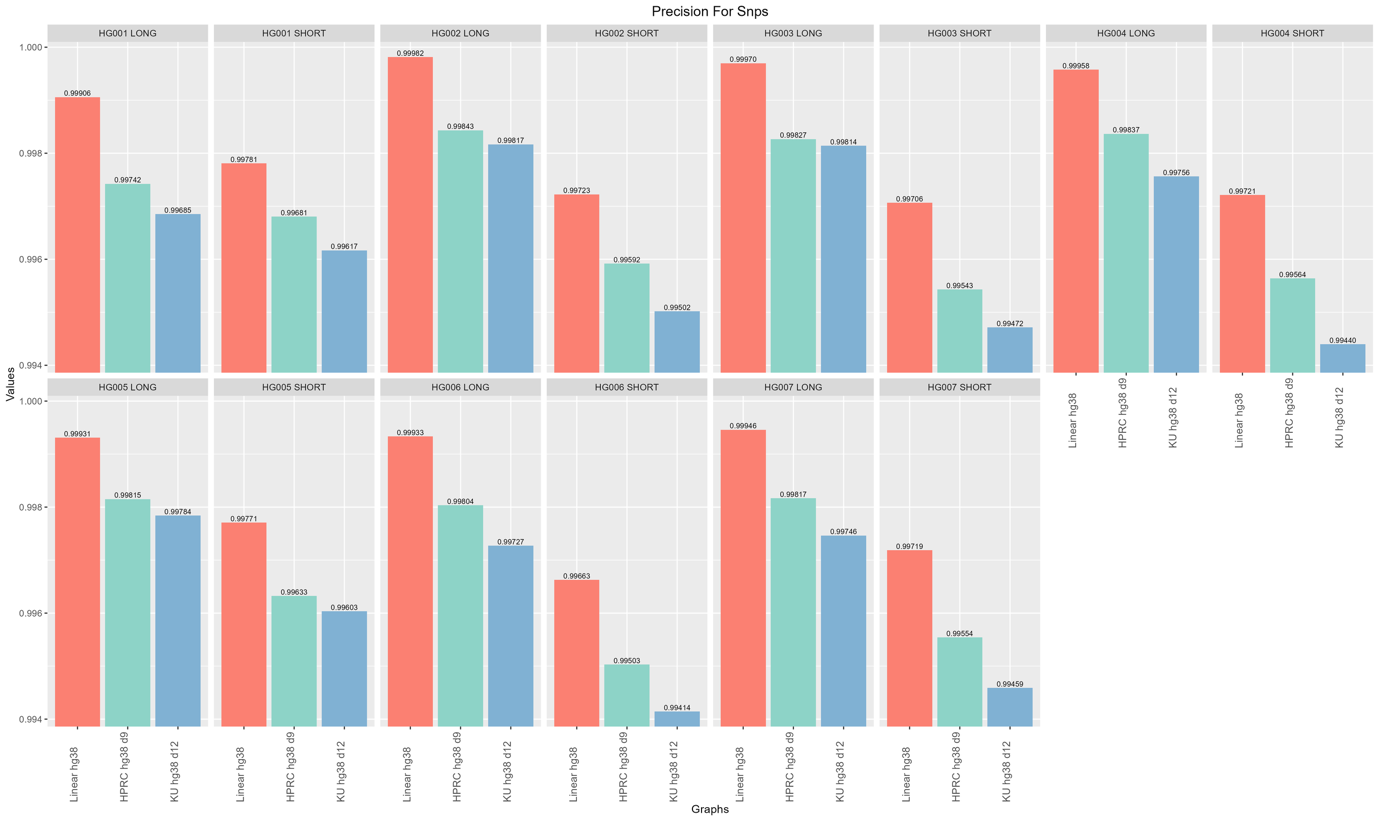

Supplementary Figure 15: Precision of SNP calls across reference genomes and sequencing platforms for GIAB samples HG001–HG007. This figure presents SNP precision metrics for Illumina (short-read) and PacBio (long-read) alignments to three reference genomes: the linear hg38, the public HPRC graph (HPRC_hg38_d9), and the Emirati T2T pangenome graph (ku_hg38_d12). SNP precision was consistently highest when aligning to the linear hg38 reference, with values reaching up to 0.9998 for long reads and 0.9978 for short reads. Graph-based references showed slightly lower precision, with long-read SNP precision ranging from 0.9968 to 0.9984, and short-read precision ranging from 0.9941 to 0.9968. Differences between HPRC and Emirati T2T pangenome graphs were minimal, with HPRC showing a marginally higher precision in most cases. The results suggest that while graph-based references perform comparably, the linear hg38 retains a consistent precision advantage for SNP calling.

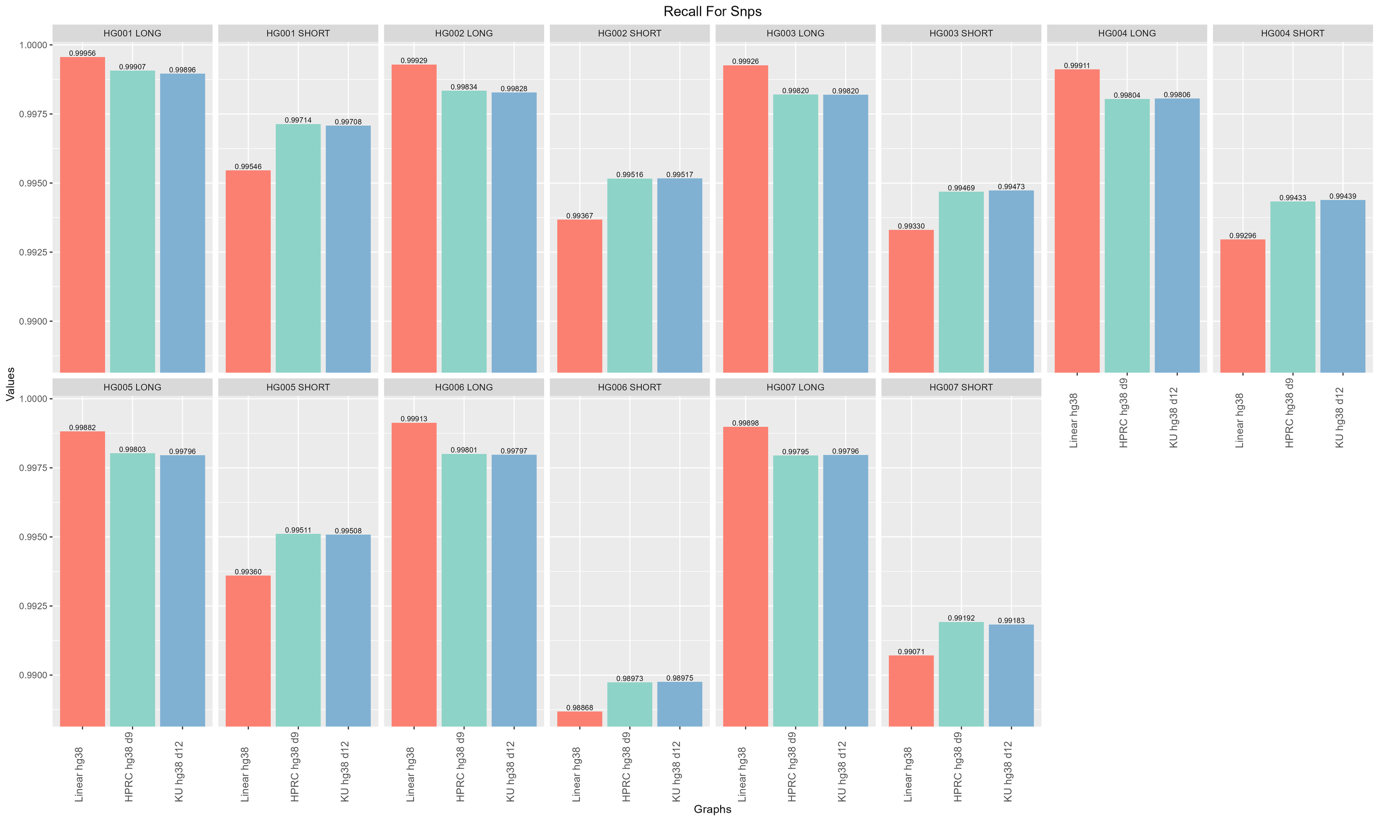

Supplementary Figure 16: SNP Recall across reference genomes and sequencing platforms for GIAB samples HG001–HG007. SNP recall values were evaluated for short-read (Illumina) and long-read (PacBio) datasets across three reference genomes: the public HPRC graph (HPRC_hg38_d9), the Emirati T2T pangenome graph (ku_hg38_d12), and the linear hg38 reference. All references demonstrated high SNP recall, with values consistently exceeding 0.988 across all samples and platforms. The linear hg38 reference achieved the highest SNP recall across most samples, particularly for long-read alignments, where recall values frequently exceeded 0.999. Differences among references were more subtle for short reads, with slightly lower recall values overall. Graph-based references (HPRC and Emirati T2T pangenome) performed comparably to each other, often trailing the linear reference by a small margin. These findings confirm high concordance in SNP detection across references but suggest a modest recall advantage for linear hg38, especially in long-read data.

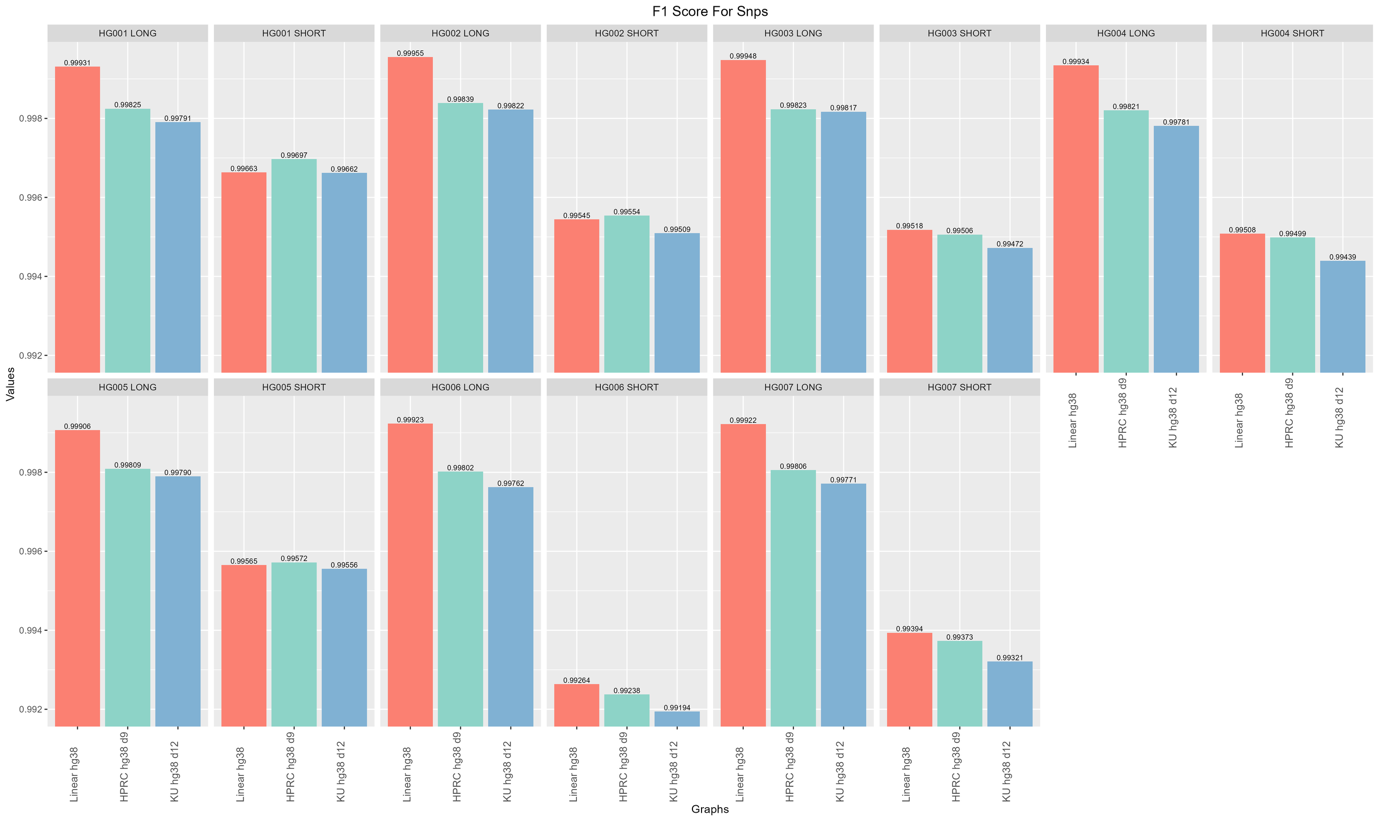

Supplementary Figure 17: F1 scores for SNP calls across reference genomes and sequencing technologies for GIAB samples HG001–HG007. This figure displays F1 scores for SNP calls from long- and short-read alignments to three reference types: the linear hg38 reference, the public HPRC graph (HPRC_hg38_d9), and the Emirati T2T pangenome graph (ku_hg38_d12). Across all samples and technologies, the linear hg38 consistently achieves the highest SNP F1 scores, with values exceeding 0.9993 for long reads and averaging 0.9955–0.9966 for short reads. HPRC and Emirati T2T pangenome graphs perform comparably but slightly lower, particularly for short reads, where F1 scores range from 0.9919 to 0.9969. Long-read SNP calls show less variability and higher consistency across references, with F1 scores above 0.9976 in nearly all cases. These results underscore the continued advantage of hg38 for SNP calling in precision-recall-balanced performance, while graph-based references approach parity but exhibit slightly diminished performance in short-read contexts.

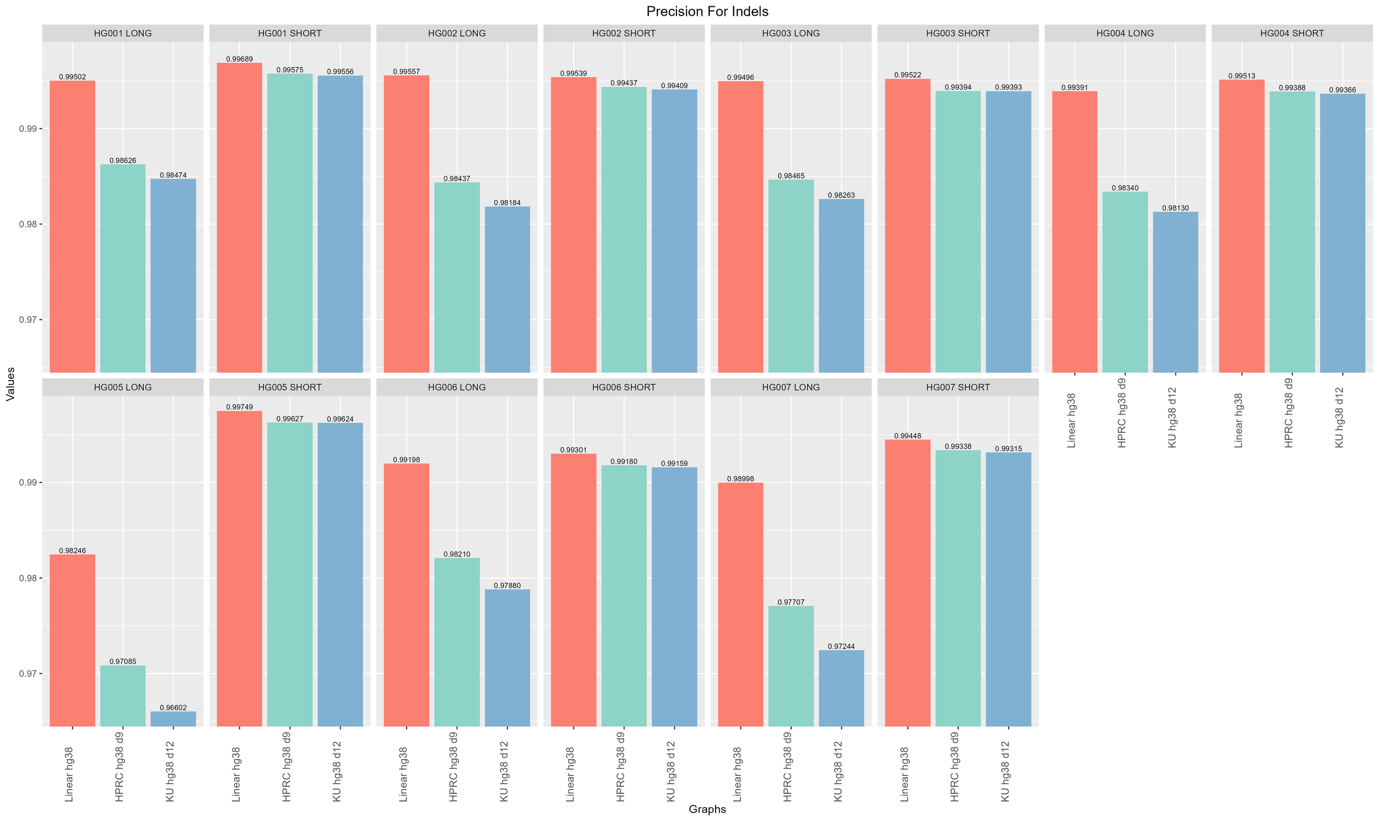

Supplementary Figure 18: Precision of indel calls across reference genomes and sequencing platforms for GIAB samples HG001–HG007. indel precision is shown for both long-read and short-read data, with alignments to the linear hg38, HPRC_hg38_d9, and ku_hg38_d12 references. Linear hg38 consistently outperformed the graph-based references, particularly for long reads, where indel precision reached up to 0.996 and showed a 1–2% advantage over HPRC and the Emirati T2T pangenome graph . Short-read indel calls demonstrated more consistent performance across references; however, hg38 still led with values of up to 0.9975, compared to 0.991–0.996 for graph-based references. The Emirati T2T pangenome graph generally trailed HPRC in long-read precision, while both performed comparably for short reads. These results underscore the continued superiority of the linear reference in indel calling, especially when using long-read technologies.

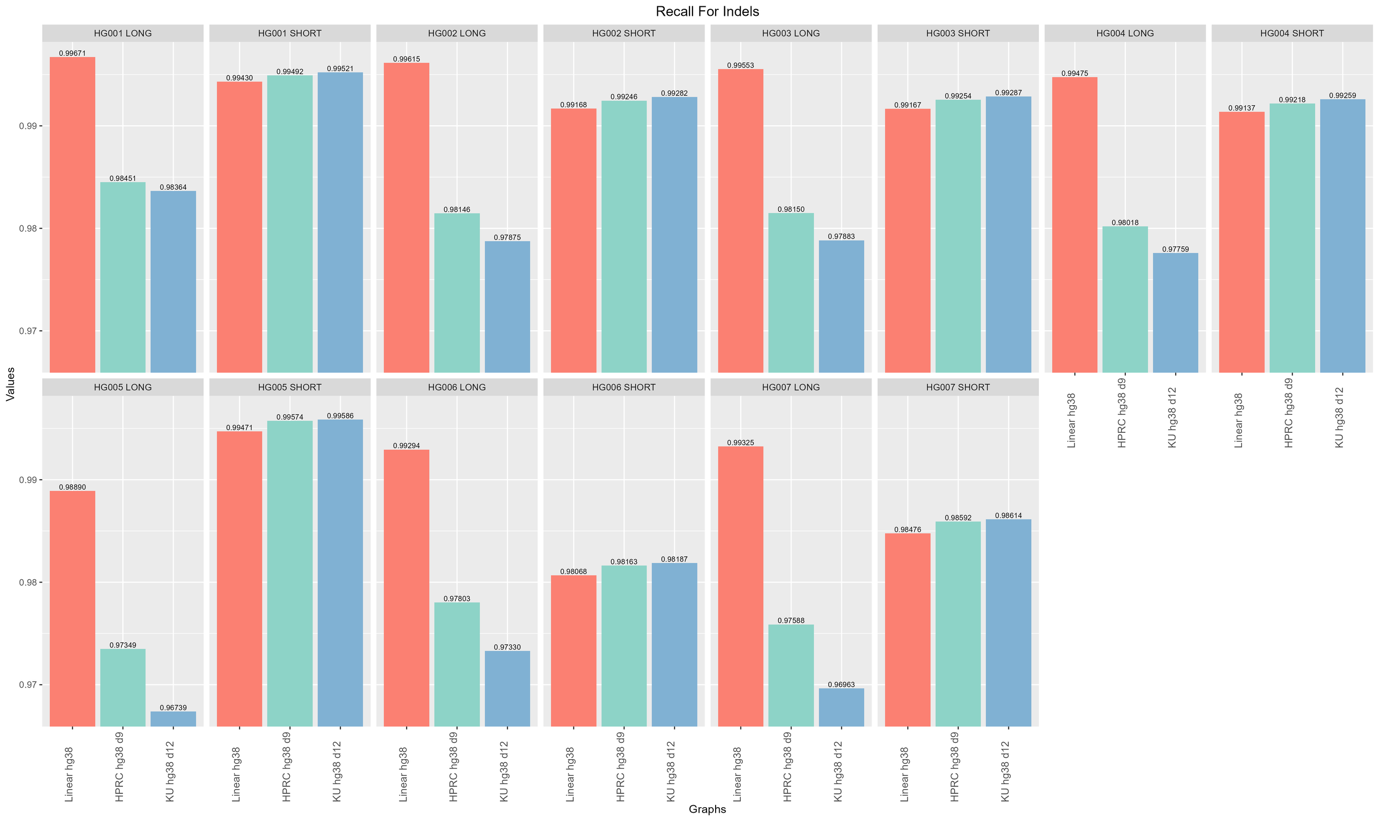

Supplementary Figure 19: Indel recall across reference genomes and sequencing platforms for GIAB samples HG001–HG007. Indel recall values were compared across Illumina and PacBio data aligned to HPRC_hg38_d9, ku_hg38_d12, and linear hg38. In contrast to SNP results, more pronounced differences emerged between references, particularly for long-read alignments. The linear hg38 reference consistently yielded the highest indel recall, often exceeding 0.99. In contrast, graph-based references (HPRC and Emirati) exhibited lower recall, particularly with long reads, with several values below 0.98 and, in some cases, near 0.96 or lower. Short-read indel recall was generally higher across all references and more uniform, with recall values approaching or exceeding 0.99. The performance gap between hg38 and the graph-based references was more evident in long-read data, suggesting potential limitations in indel detection or alignment specificity when using current graph-based tools for this variant class.

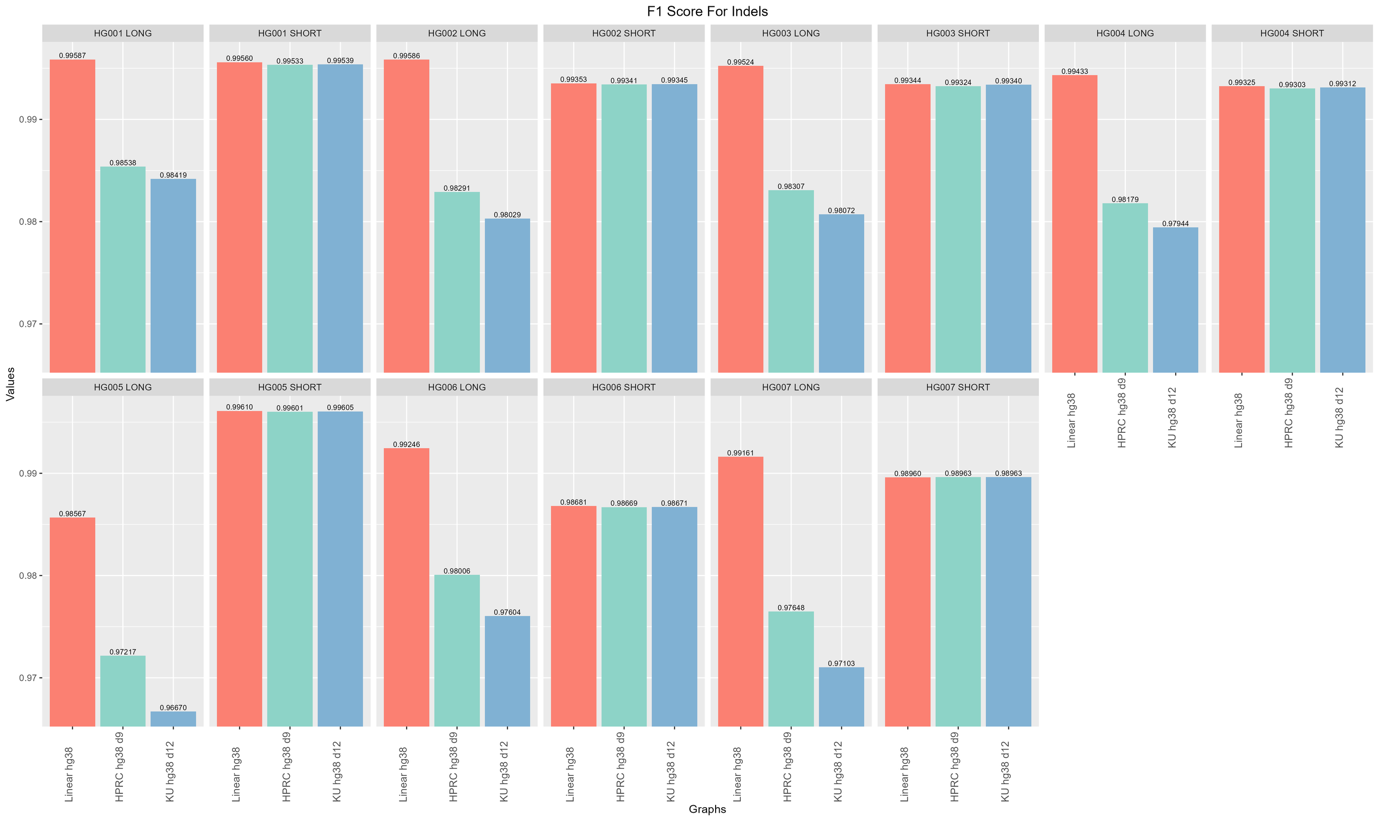

Supplementary Figure 20: F1 scores for indel calls across reference genomes and sequencing technologies for GIAB samples HG001–HG007. This figure summarizes indel F1 scores for short- and long-read alignments to the linear hg38, HPRC_hg38_d9, and ku_hg38_d12 references. The linear hg38 reference consistently yields the highest indel F1 scores, reaching up to 0.9961 for short reads and 0.9959 for long reads. In contrast, graph-based references exhibit a performance drop, particularly for long-read inputs, where HPRC and Emirati T2T pangenome F1 scores can fall below 0.98, and in some samples, below 0.97. For short reads, the differences are more modest, with F1 scores for all references clustering tightly between 0.986 and 0.996. These data highlight the relative robustness of the linear reference in indel detection, especially for long-read sequencing, and suggest further optimization is needed for graph-based approaches in complex variant contexts.

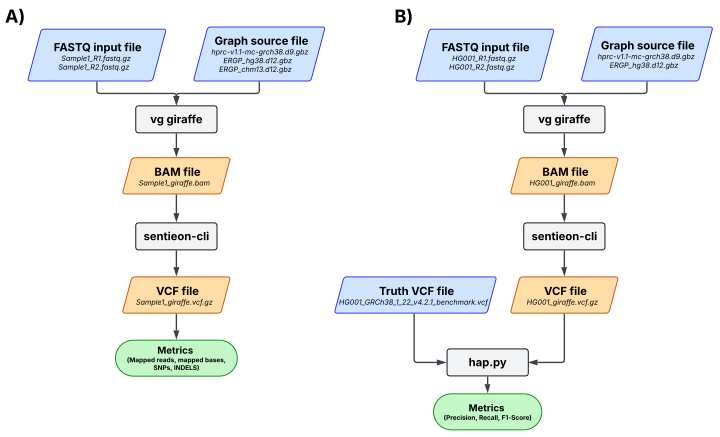

Supplementary Figure 21: Variant-calling workflows for short-read data. (A) Novaseq 6000 short-read paired-end FASTQs (Sample1_R1/R2) for 119 in-house Emirati samples are aligned to population graph references (HPRC v1.1 multi-assembly GRCh38 and Emirati T2T pangenome graphs for hg38/CHM13) using vg giraffe, variants are then called with sentieon-cli to yield compressed VCFs, and run-level metrics (mapped reads/bases, SNV/indel counts) are collected. (B) Truth-set samples (HG001-HG007) are processed with the same alignment and calling steps; resulting VCFs are assessed against the GIAB truth VCF (HGxxx_GRCh38_1_22_v4.2.1) using hap.py to report precision, recall, and F1-score.
